# Proteomics of broad deubiquitylase inhibition unmasks redundant enzyme function to reveal substrates

**DOI:** 10.1101/844811

**Authors:** Valentina Rossio, Joao A. Paulo, Joel Chick, Bradley Brasher, Steven P. Gygi, Randall W. King

**Author notes:** Correspondence to Randall W. King.

## Abstract

Deubiquitylating enzymes (DUBs) counteract ubiquitylation to control stability or activity of substrates. Identification of DUB substrates is challenging because multiple DUBs act on the same substrates, thwarting genetic approaches. Here, we circumvented redundancy by broadly inhibiting DUBs in *Xenopus* egg extract. DUB inhibition increases ubiquitylation of hundreds of proteins, depleting free ubiquitin without inducing widespread degradation. Restoring available ubiquitin led to proteasomal degradation of over thirty proteins, indicating that deubiquitylation is essential to maintain their stability. We confirmed their DUB-dependent stability with recombinant human proteins, demonstrating evolutionary conservation. We profiled the ability of DUBs to rescue protein stability, and found that USP7 has a unique ability to broadly antagonize proteasomal degradation. Together, we provide a comprehensive characterization of ubiquitin dynamics in the *Xenopus* system, identify new DUB substrates, and present a new approach to characterize physiological DUB specificity that overcomes challenges posed by DUB redundancy

## Introduction

Conjugation of ubiquitin to proteins is a widespread, highly regulated post-translational modification. The Ubiquitin-Proteasome System (UPS) is best known for its ability to target specific proteins for degradation. Still, ubiquitylation also regulates protein localization, activity and function independently of degradation(Swatek and Komander 2016),(Hershko and Ciechanover A. 1998),(Clague, Heride, and Urbé 2015). Ubiquitin can be incorporated in polyubiquitin chains of different topologies that may result in different fates(Komander and Rape 2012). For example, lysine-63 linked chains promote non-degradative functions (Jackson and Durocher 2013),(Lauwers, Jacob, and André 2009), whereas lysine-48 linked chains and branched chains such as K11/K48 or K29/K48 promote proteasomal degradation(Chau et al. 1989). Ubiquitin is covalently attached to substrates via a cascade of E1-E2-E3 enzymes(Yihong Ye and Rape 2009). In contrast, deubiquitylating enzymes (DUBs) catalyze ubiquitin removal(Clague, Coulson, and Urbé 2012).

Comprehensive proteomic identification of physiological UPS substrates has been challenging due to the prominence of protein-quality control pathways that ubiquitylate newly synthesized proteins that do not fold properly in growing cells(W. Kim et al. 2011). Therefore, it can be difficult to distinguish regulatory ubiquitylation of mature proteins from more widespread ubiquitylation that targets misfolded proteins for degradation during protein biogenesis. To overcome this challenge, in our study we took advantage of the *Xenopus laevis* egg extract model system. Since the mature *Xenopus* egg has completed its growth prior to being laid, there is little ongoing translation and protein folding, thereby minimizing the contribution of quality-control ubiquitylation compared to actively growing cultured cells. Thus, ubiquitylation events observed in these extracts are more likely to reflect physiologic regulatory events rather than quality control.

*Xenopus* egg extract can be prepared with little dilution of the cytoplasm, therefore preserving physiological integrity and enabling the reconstitution of complex biochemical processes such as DNA damage repair cell cycle progression and mitotic spindle formation (Yardimci et al. 2012),(Glotzer, Murray, and Kirschner 1991),(Field and Mitchison 2018). *Xenopus* egg extract has been routinely used for studies of protein ubiquitylation and degradation, including studies of mitotic cyclin degradation(Glotzer, Murray, and Kirschner 1991) as well as regulatory ubiquitylation during DNA replication(Yardimci et al. 2012). Together these studies indicate that ubiquitylation regulates important physiological processes in *Xenopus* egg extracts. Still, we have a limited understanding of ubiquitin homeostasis and the overall dynamics of ubiquitylation, deubiquitylation, and protein degradation in this experimental system.

The human genome encodes approximately 100 DUBs, divided into two families: the zinc metalloprotease class (10 DUBs), and the cysteine-protease class that contains most other DUBs (~90)(Mevissen and Komander 2017),(Clague et al. 2013). A few DUBs have been well-studied with many characterized substrates, but most DUBs still do not have any known substrates, limiting our understanding of how DUB substrate specificity is achieved. DUB specificity is complex, as it can arise through binding specificity mediated by unique protein-protein interaction domains(Ma et al. 2010), or through recognition of specific topologies of ubiquitin chains(Bremm, Freund, and Komander 2010; Wang et al. 2009),(Flierman et al. 2016). Most DUB substrates have been discovered by first identifying proteins that interact with specific DUBs. This approach has been used globally(Sowa et al. 2009), but it may miss substrates that interact weakly with DUBs or identify regulators or scaffold proteins rather than true substrates. Identifying DUB targets is difficult because DUBs may function redundantly(Beckley et al. 2015),(Kwon, Saindane, and Baek 2017), and thus inactivating a single DUB may not destabilize its substrates or affect a particular biological process. In the case of proteome stability, it is not known which DUBs have the broadest impact in rescuing proteins from proteasome degradation or to what extent DUBs act redundantly in this process. Therefore, new approaches are needed to identify physiological substrates of DUBs and to overcome the challenges posed by redundant function of these enzymes.

In this study we applied unbiased quantitative proteomic approaches to characterize the dynamics of ubiquitylation and proteasomal degradation in *Xenopus* egg extract. We found that the proteome is stable, despite widespread ubiquitylation, suggesting that most ubiquitylation is non-degradative. By broadly inhibiting cysteine-protease DUBs, we circumvented the possible effects of redundancy on proteome stability and discovered a set of proteins whose stability depends on DUB activity. We next took advantage of this panel of substrates to identify DUBs that are sufficient to counteract proteasomal degradation of these proteins. By broadly inactivating DUBs and adding back a panel of recombinant DUBs one by one, we unmasked DUB redundancy and discovered that USP7 can rescue many substrates from degradation. However, specific inhibition of USP7 with a small molecule inhibitor was not sufficient to promote degradation of most of the substrates we identified, suggesting that USP7 functions redundantly with other DUBs. Our work highlights the impact of DUB redundancy on proteome stability and reveals the specificity and activity of DUBs whose function would otherwise be masked by redundancy.

## Results

### UbVS treatment induces rapid depletion of available ubiquitin and labels a broad set of cysteine-protease DUBs in *Xenopus* extract

To selectively inhibit cysteine protease DUBs in Xenopus extract, we used ubiquitin vinyl sulfone (UbVS), which covalently inhibits cysteine-protease DUBs, without inhibiting other cysteine proteases or other classes of DUBs(Borodovsky et al. 2001). Our laboratory previously showed that UbVS treatment efficiently blocks ubiquitin recycling in *Xenopus* extract(Dimova et al. 2012). We confirmed that 10 μM UbVS was sufficient to rapidly deplete free ubiquitin, (Fig. 1a, 1b) which was associated with rapid discharge of the Ube2C-ubiquitin thioester within 5 minutes (Fig. 1a, bottom). Other E2s were similarly rapidly discharged (data not shown). Addition of exogenous ubiquitin to UbVS-treated extract fully recharged the Ube2C thioester confirming that the E2’s discharge was due to the depletion of free ubiquitin (Fig.1a, bottom). To determine the spectrum of cysteine-containing DUBs targeted by this concentration of UbVS, extract was treated with 10 μM HA-tagged UbVS (HA-UbVS) and sensitive DUBs were visualized as discrete bands by anti-HA Western blotting analysis (Fig. 1b, right). To identify these DUBs, we performed mass spectrometry analysis of immunopurified HA-UbVS from extract and identified 88 proteins (Table S2), 35 of which were cysteine-protease DUBs (Table S1, S2) that are likely direct targets of UbVS. All known classes of cysteine-protease DUBs were found (Fig. 1c, Table S1), whereas metallo-protease DUBs (JAMM) were not identified, as expected(Borodovsky et al. 2001). Prior proteomic analysis of *Xenopus* extract detected 54 DUBs(Wühr et al. 2014) (Table S1), 51 of which were cysteine-proteases. Thus, 10 μM HA-UbVS labeled 35 of 51 (69%) of cysteine-protease DUBs present in extract, consistent with broad specificity of UbVS for this class of DUBs(Borodovsky et al. 2001). The remaining DUBs were likely not identified either because they were present in low abundance or because they do not react rapidly with UbVS. Our HA-UbVS pull-down also isolated proteins that are likely to strongly associate with DUBs (Table S2), as the pull-down was carried out in the presence of high salt (500 mM KCl). A high fraction of these proteins (31) are components of the proteasome, consistent with the known interaction of UbVS-sensitive DUBs UCHL5 and USP14(de Poot, Tian, and Finley 2017) with the proteasome. Other known interactors of DUBs that we isolated included GBP1/GBP2, which binds USP10(Soncini, Berdo, and Draetta 2001), and the SPATA proteins (SPATA2/SPATA2L), which have been identified as CYLD interactors(Sowa et al. 2009) in human cells (Table S2), suggesting evolutionary conservation of these interactions.

**Fig.1.**
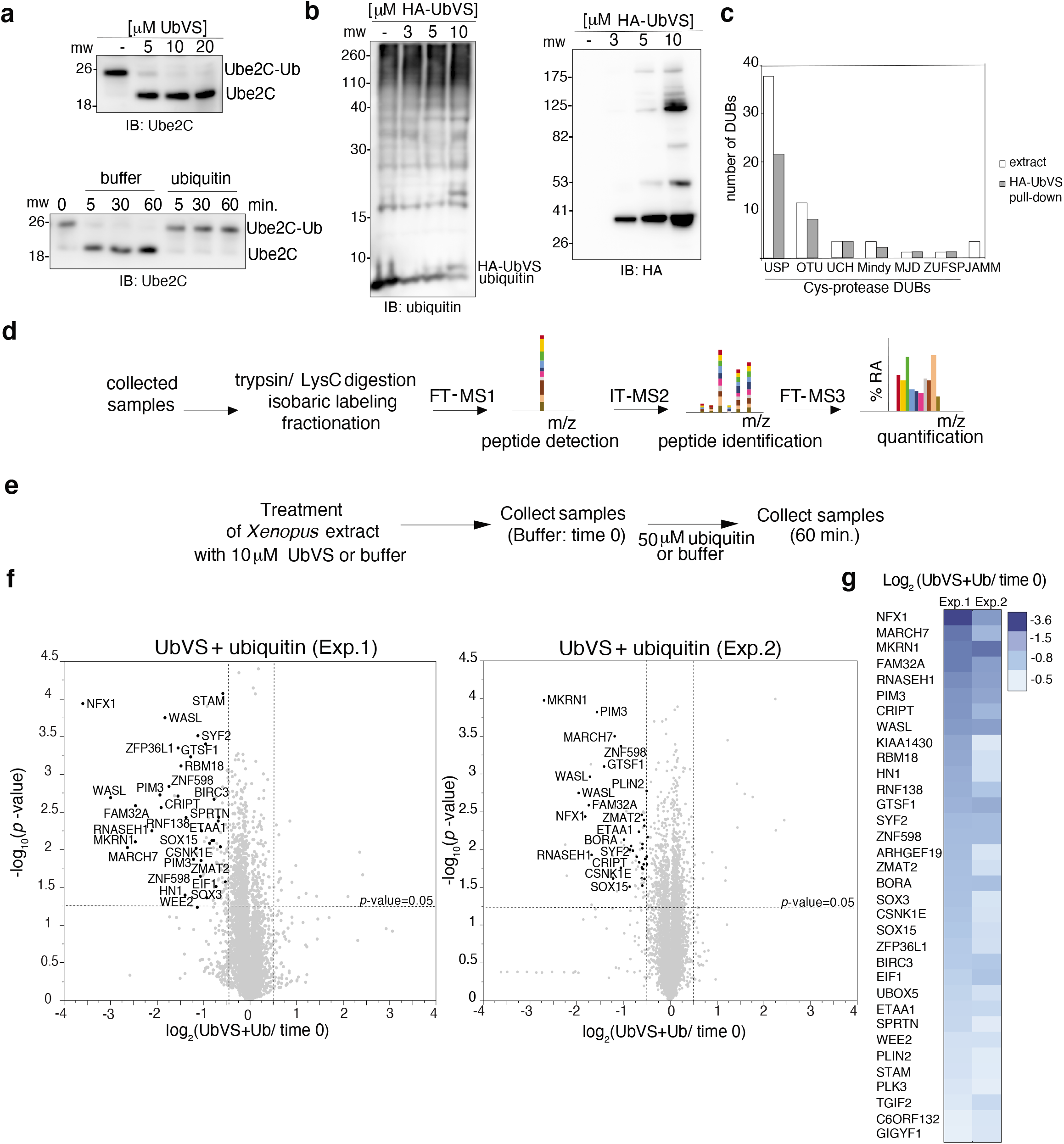
Inhibition of cysteine-protease DUBs by UbVS causes rapid ubiquitin depletion and degradation of a set of proteins. **(a)** Extract was treated with UbVS or **(b)** HA-UbVS. After 10 minutes (time 0), 50 μM ubiquitin or buffer were added (**a**, bottom). Samples were subjected to immunoblotting (IB). **(c)** Extract was incubated with 10 μM HA-UbVS (10 minutes) and subjected to anti-HA pull-down followed by mass spectrometry analysis.The number of DUBs identified (gray) is compared to the number of DUBs detected previously in extract (Wuhr et al. 2014) (white). **(d)** Mass spectrometry analysis by SL-TMT method is summarized **(e)** Ubiquitin or buffer were added to the extract (time 0) pre-incubated with buffer or UbVS (10 minutes). Samples were collected as indicated **(f)** Volcano plots of TMT analysis comparing the proteins detected in UbVS/ubiquitin at 60 minutes with the proteins detected at time 0 (2 independent experiments). Statistical significance (−log_10_ *p-value*) is plotted against the log-transformed ratio of the samples. Proteins significantly decreasing in both the experiments are shown in black. **(g)** Heat map of the proteins downregulated in UbVS/ubiquitin in both the experiments in **(f).**

### UbVS treatment induces degradation of a limited set of proteins when available ubiquitin is restored

We first wanted to investigate how inhibition of multiple DUBs influenced proteome stability in *Xenopus* extract. We hypothesized that simultaneous inhibition of 35 DUBs with UbVS might lead to destabilization of proteins that require ongoing deubiquitylation to maintain their stability. By simultaneously inhibiting a large number of DUBs, we predicted that we might be able to identify new substrates of DUBs, including substrates whose stability is conferred by the action of redundant DUBs. We used a multiplexed Tandem Mass Tag (TMT)-based quantitative proteomic approach(Navarrete-Perea et al. 2018) (Fig. 1d), and we measured protein abundance over time in untreated extract, as well as extract to which ubiquitin, UbVS, or the combination (UbVS/ubiquitin) were added (Fig. 1e). We reasoned that proteins that decreased specifically in the presence of UbVS and/or UbVS/ubiquitin may normally be protected from degradation by UbVS-sensitive DUBs. We performed two independent experiments using *Xenopus* extract prepared from two sets of eggs (collected from different animals and processed separately), to ensure reproducibility. In each experiment we quantified around 8000 proteins, with an overlap of 94.8% between the two experiments.

First, we observed that in the absence of any perturbation, the proteome was stable in *Xenopus* extract over the course of 60 minutes. Few proteins changed in abundance over time, and those that did were not shared between the experiments (Fig. S1a). This degree of proteome stability is consistent with the idea that bulk rates of translation are relatively quiescent in the *Xenopus* system(Richter and Smith n.d.) and that the extract was treated with cycloheximide to prevent protein synthesis and mitotic entry. Proteome stability in untreated extract was not a consequence of limiting ubiquitin availability because addition of exogenous ubiquitin alone did not stimulate protein degradation (Fig. S1a). With the exception of ubiquitin, which was added to the extract, we did not observe any proteins that increased in abundance (Fig. S1a) in both the experiments. Addition of UbVS alone also had no measurable effect on protein stability, possibly due to depletion of available ubiquitin (Fig. S1a). However, addition of UbVS together with ubiquitin (UbVS/ubiquitin) induced degradation of 34 proteins, as defined by a reduction in their abundance by at least 1.5-fold in both experiments (Fig. 1f, g). Choosing a more relaxed threshold based on the 5% of proteins whose abundance decreased most in UbVS/ubiquitin (FDR 1%) revealed additional proteins that decreased in both experiments (Fig. S4a). Because transcription and translation were inactive in extract, the decrease in protein abundance of this specific set of proteins was likely a consequence of protein degradation that occurred as a direct consequence of DUB inhibition. Thus, these proteins represent putative DUB substrates that are protected from degradation by UbVS-sensitive DUBs.

Because only a limited number of proteins were destabilized by the addition of 10 μM UbVS/ubiquitin, we tested whether stronger inhibition of UbVS-sensitive DUBs, by addition of a higher concentration of UbVS, led to destabilization of a larger number of proteins. We compared protein stability in untreated extract and in extract co-treated with ubiquitin and 10 μM (as in the previous experiments) or 30 μM UbVS (Fig. S2a) using TMT-based quantitative proteomics as before. We confirmed the degradation of the proteins observed previously using 10 μM UbVS, but we did not observe an increase in the number of proteins degraded in the presence of 30 μM UbVS (Fig. S2b). On the contrary, the proteins decreasing in 10 μM UbVS were more stable at the higher UbVS concentration (Fig. S2b, S2c). This finding suggested that addition of a higher concentration of UbVS might lead to faster ubiquitin depletion, thereby hampering the ability of UbVS to stimulate protein degradation. To test this hypothesis, we monitored the discharge rate of the E2-Ub thioester after ubiquitin addition to extract pre-treated with increasing concentrations of UbVS. Whereas addition of 50 μM ubiquitin was sufficient to completely sustain Ube2C charging for 30 minutes in extract treated with 10 μM UbVS, this concentration of ubiquitin was insufficient to maintain charged Ube2C in extract treated with 30 μM UbVS (Fig. S2d). Thus, we cannot drive broader protein instability by stronger DUB inhibition because ubiquitin becomes depleted too rapidly.

### Non-degradative ubiquitylation targets a large number of proteins in *Xenopus* egg extract

Since addition of UbVS causes rapid depletion of available ubiquitin and discharge of E2 enzymes (Fig.1a, b), the ubiquitin conjugation machinery must be active in *Xenopus* extract. We were therefore surprised by the fact that addition of ubiquitin or UbVS alone did not stimulate protein degradation (Fig.S1a) and that UbVS/ubiquitin promoted degradation of only a limited set of proteins (Fig.1g). These findings may be explained by the prevalence of non-degradative ubiquitin conjugation pathways that preferentially consume ubiquitin, limiting ubiquitylation of degradative substrates. To acquire a better picture of the spectrum of proteins that are ubiquitylated in extract, we performed a proteomic experiment to identify proteins that become modified with exogenously added ubiquitin. We treated the extract with 50 μM of HA-ubiquitin or buffer for 30 minutes to allow the incorporation of HA-ubiquitin into ubiquitylated substrates. Immunoblot revealed incorporation of tagged ubiquitin into many proteins (Fig. S3a). We then performed mass spectrometry analysis of immunopurified HA-ubiquitin from extract and identified 772 proteins that were enriched for binding to HA-antibody beads relative to empty beads (log2 anti-HA beads relative to empty beads)> 1, p-value < 0.05) (Fig.S3b). Because this pull-down was carried out in the presence of high salt (500 mM KCl), these proteins are likely either directly ubiquitylated or bind ubiquitin with high affinity. The top proteins enriched on the anti-HA beads were UPS components such as E1s, E2s and ubiquitin ligases (Fig. S3B) that can form a thioester with HA-ubiquitin, confirming the validity of this approach. Beyond ubiquitylation machinery, proteins involved in translation, such as ribosomes, translation factors and RNA binding proteins, were the most frequently identified. Other proteins isolated included protein folding factors, cytoskeletal components, proteins involved in DNA replication/repair, mitochondrial proteins, and metabolic enzymes. We also identified 9 of the 34 proteins degraded in extract treated with UbVS/ubiquitin, confirming that these proteins are directly ubiquitylated in extract. Thus, while the proteome is stable in untreated extract, the ubiquitin conjugation machinery appears highly active in its ability to modify a large number of proteins with ubiquitin in a manner that does not impact their stability. Therefore, the failure of ubiquitin addition to stimulate widespread degradation (Fig.S1a) may be a result of the preferential incorporation into non-degrative substrates.

Together our findings suggest that UbVS treatment induces rapid ubiquitin depletion principally by promoting ubiquitylation of non-degradative substrates. To identify proteins that become preferentially ubiquitylated in the presence of UbVS, we profiled global protein ubiquitylation in extract using a multiplexed quantitative di-glycine (diGly) remnant method(Rose et al. 2016) after addition of ubiquitin or UbVS/ubiquitin to the extract for 30 minutes. Overall, we identified 883 ubiquitylation sites in 515 proteins, 219 of which were found in the previous HA-ubiquitin pull down experiment (Fig. S3c), indicating substantial overlap with the two different approaches. We identified 190 ubiquitylation sites (142 proteins) whose ubiquitylation increased significantly after UbVS/ubiquitin addition compared to addition of ubiquitin alone (log_2_ Fold change>1) (Fig.S3d). As expected, we found 8 ubiquitylation sites belonging to 6 proteins of the 34 that were destabilized by UbVS/ubiquitin addition to the extract (Fig. S3e), confirming again that these proteins are direct DUB substrates. Several proteins identified have been previously reported to be DUB substrates, such as the ESCRT complex component CHMP1B (USP8)(Crespo-Yàñez et al. 2018), the ubiquitin ligase SMURF2 (USP15)(Iyengar et al. 2015) and the replication factor PCNA (USP1)(Huang et al. 2006). However, the majority of the UbVS-sensitive sites are novel. Because UbVS treatment increases ubiquitylation of a much larger number of proteins than those that are destabilized, including abundant proteins such ribosome subunits and histones, these substrates may sequester ubiquitin, limiting protein degradation in extract treated with UbVS alone.

### Confirmation of UbVS-dependent proteasomal degradation of the newly identified substrates with human orthologs

After developing a clearer picture of the overall dynamics of ubiquitylation, deubiquitylation, and protein degradation in extract, we next focused on the 34 substrates that require UbVS-sensitive DUBs to maintain their stability. Most of these proteins have reported physiological functions, but only 9 are known UPS substrates (Fig. 2a, Fig. S4a, Table S3). Furthermore, for only four of them have specific DUBs been identified that control their stability. Two of these proteins are ubiquitin ligases, MARCH7 and BIRC3, which are known to be protected from degradation by cysteine-protease DUBs in human cells (by USP7 or USP9(Nathan et al. 2008) and USP19 respectively(Mei et al. 2011)). In addition, we identified Stam and NFX1, which are known to be deubiquitylated by the cysteine-protease DUBs USP8(Berlin, Schwartz, and Nash 2010) and USP9 respectively(Chen et al. 2019). The ability of our screen to identify known DUB substrates suggests that our approach is capable of identifying physiologically relevant proteins whose stability requires DUB activity.

**Fig. 2.**
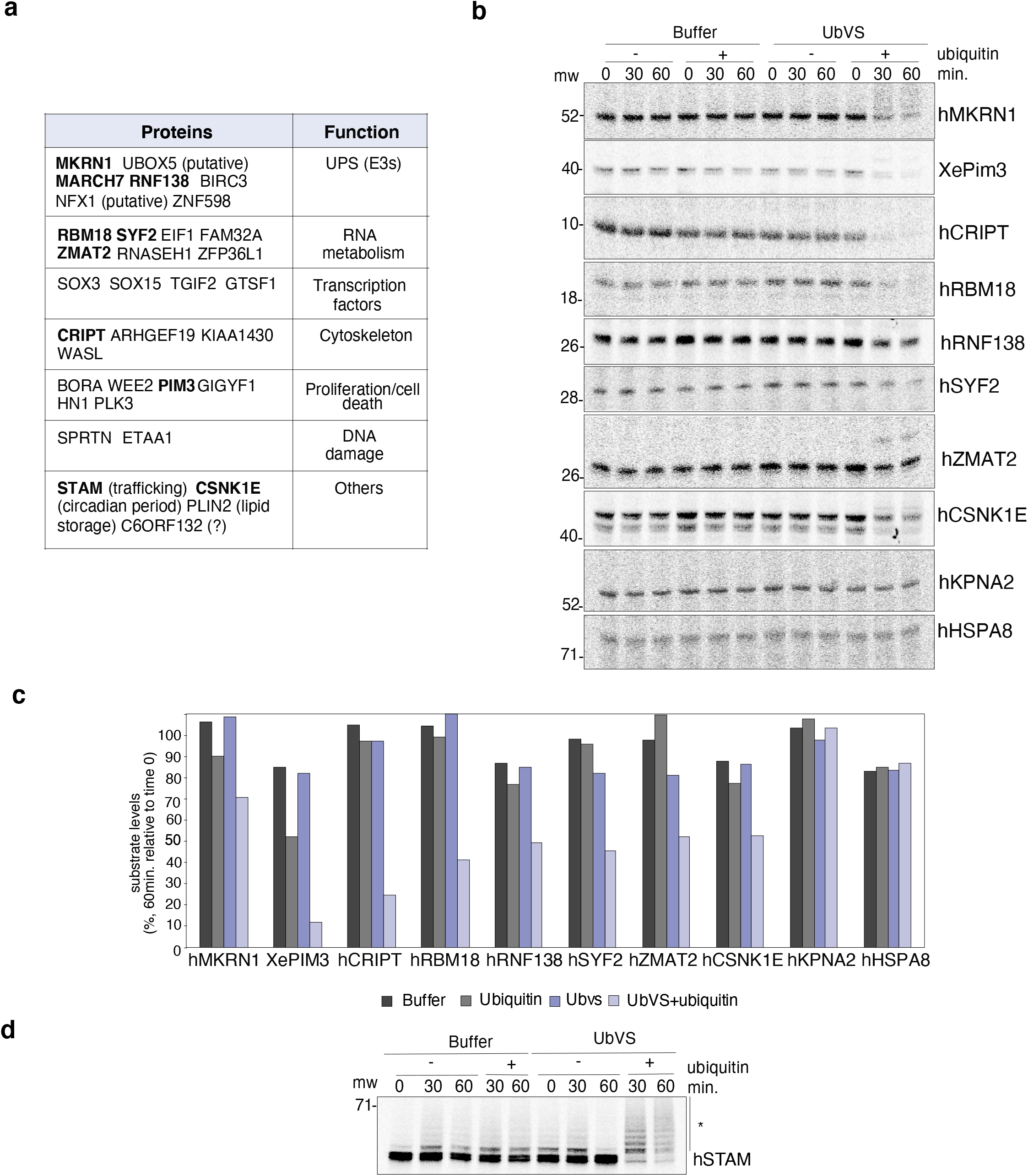
Independent validation of the proteins protected from degradation by UbVS-sensitive DUBs. **(a)** Functions of the proteins down-regulated in UbVS/ubiquitin-treated extract (Fig.1g). Proteins independently validated are in bold. **(b)** Human orthologs (prefix h) or Xenopus protein (prefix Xe) were expressed in reticulocyte lysate, labeled with 35S-methionine and added to the extract, treated as in Fig.1e. Aliquots were collected and processed for SDS-PAGE and phosphorimaging. UbVS/ubiquitin-dependent degradation of these proteins was confirmed in 2 independent experiments. One of the expeiments is shown **(c)** Quantification of the experiment in **(b).** Quantification of the second experiment is shown in Fig. S3d **(d)**Performed as in **(b)** *: ubiquitylated species.

For the majority of the proteins we identified, DUBs have not been implicated in regulation of their stability. We identified four additional ubiquitin ligases (MKRN1, RNF138, ZNF598, UBOX5), whose stability is not known to be dependent on DUB activity. This enrichment for ubiquitin ligases is again consistent with the fact that DUBs are known to protect them from auto-ubiquitylation and consequent degradation(W. Kim et al. 2011),(Ventii and Wilkinson 2008). Other functional classes of proteins that we identified included transcriptional regulators (GTSF1, SOX3, SOX15, TGIF2), signaling proteins (GYGIf1, PLK3, WEE2, CSNK1E, BORA, PIM3), cytoskeletal regulators (CRIPT, WASL, KIAA1430, HN1, ARHGEF19), proteins involved in RNA processing (FAM32, RNASEH1, RBM18, SYF2, ZMAT22, ZFP36L11, EIF1), DNA damage components (ETAA1, SPRTN), a lipid storage protein (PLIN2), and an uncharacterized protein (C6ORF132). Together these findings suggest that ongoing deubiquitylation in the *Xenopus* system is important for maintaining the stability of proteins that regulate a wide variety of biological processes.

We looked for possible characteristics sequence of these substrates (Table S3) that could contribute to their recognition by the UPS. Consistent with the PEST sequence(Rechsteiner and Rogers 1996) being a signal that promotes ubiquitin and proteasome-dependent degradation, we found one or multiple PEST sequences in 13 of the 34 hits (Table S3). Because efficient protein degradation by the proteasome requires unstructured regions in its substrates(Aufderheide et al. 2015), we analyzed their sequences using ProViz(Jehl et al. 2016). We found that 8 substrates were predicted to be completely or highly disordered whereas 25 had significantly disordered regions consistent with the possibility that they are proteasome substrates (Table S3).

To directly test whether degradation of the identified DUB substrates was proteasome-dependent, we quantified protein stability in extract treated with UbVS/ubiquitin in the presence of the proteasome inhibitor MG262 or DMSO (as a control) with a TMT-based quantitative proteomics experiment (Fig. S5a, S5b). We found that 19 proteins became unstable after UbVS/ubiquitin addition (Fig. S5b, S5c), 14 of which were among the 34 substrates identified (Fig. 1g). All of these proteins were stabilized by proteasome inhibition (Fig. S5b, c). Thus, UbVS-sensitive DUBs antagonize proteasome-mediated degradation of these proteins.

To verify the findings of the proteomic experiments and directly test whether these proteins become destabilized by DUB inhibition, we expressed 13 candidate substrates in rabbit reticulocyte lysate and labeled them with ^35^S-methionine. We chose ten proteins from our list of 34 candidates, as well as three proteins that were destabilized in at least one of the two experiments shown in Fig.1f. We translated *in vitro* human orthologs, with the exception of PIM3, where we tested the *Xenopus* protein. Subsequently, we added the translated proteins to untreated extract or to extract pre-treated with ubiquitin, UbVS or UbVS/ubiquitin and monitored their stability over time. We found that 12 of 13 proteins were degraded in the presence of UbVS/ubiquitin but were stable in the other conditions (Fig. 2b, 2c, S4b-d), showing the same pattern that we observed for the endogenous counterparts. STAM was the only protein that was not destabilized in the presence of UbVS/ubiquitin, but instead showed strong poly-ubiquitylation (Fig. 2d). As a control, two proteins that were stable in the proteomic experiments (KPNA2 and HSPA8), were also tested and found to be stable when assayed by this approach (Fig. 2b, 2c) Thus, because we could recapitulate the behaviour of the endogenous *Xenopus* proteins using human orthologs, the DUB-dependent regulation of these proteins is likely conserved, as is known already for MARCH7, BIRC3, NFX1 and STAM.

### Identification of DUBs sufficient to rescue proteins from degradation in UbVS/ubiquitin-treated extract

Next, we used these substrates to investigate the role of cysteine-protease DUBs in counteracting their proteasomal degradation. If the proteins degraded in UbVS/ubiquitin-treated extract are true DUB substrates, addition of recombinant DUBs to the extract should be able to rescue their degradation. Furthermore, by testing the sufficiency of each DUB to rescue degradation of these substrates, we can evaluate the activity and specificity of each DUB, even if they normally function redundantly. We reasoned that the DUBs efficiently targeted by UbVS were most likely to rescue the stability of the proteins degraded. Thus, we measured the fraction of each DUB labeled by UbVS. After we treated the extract with 10 μM HA-UbVS, we subsequently depleted all the HA-UbVS (and associated proteins) from the extract by immunodepletion with anti-HA antibodies coupled to beads. Using TMT-based quantitative proteomics, we compared the proteins remaining in extract after immunodepletion of HA-UbVS to the proteins present in extract after incubation with empty beads. As expected, the proteins depleted were mostly DUBs (Fig. S6a). We detected 32 cysteine-protease DUBs in undepleted extract and found that 25 of them were depleted by HA-UbVS (Fig. 3a), with 19 DUBs being depleted by at least 30%. There was no clear correlation between the reported abundance(Wühr et al. 2014) of the DUBs (Table S1) and their fractional depletion by HA-UbVS (Fig. 3a). We selected 12 DUBs that were depleted at least 30% to test if they could rescue substrate degradation in UbVS/ubiquitin treated extract (Fig.3a, in bold). In addition, we tested USP14 because it can rescue several substrates from proteasomal degradation(B.-H. Lee et al. 2010a), as well as USP21 because it is a highly active DUB widely used in in vitro assays.

**Fig 3.**
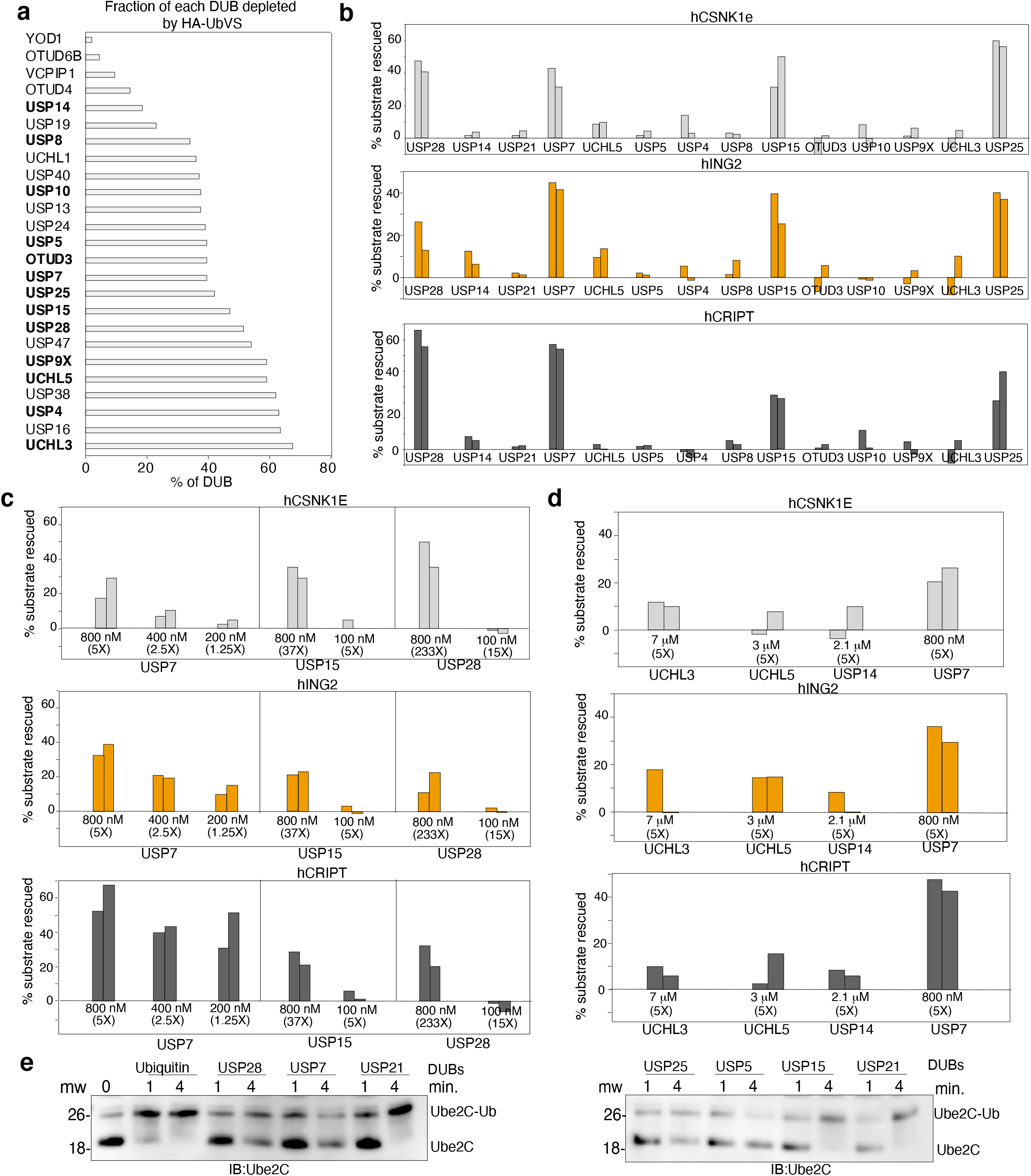
Identification of DUBs sufficient to rescue degradation of the selected substrates in UbVS/ubiquitin-treated extract. **(a)** Percentage of DUBs depleted by HA-UbVS. Extract was treated with 10μM HA-UbVS (10 minutes) and incubated overnight with anti-HA or empty beads. Supernatants were analyzed in triplicate by the SL-TMT method. Bold: DUBs tested in subsequent experiments. **(b)** Extract was incubated with 10 μM UbVS (30 minutes),after which recombinant DUBs (800 nM), labeled substrates and ubiquitin were added. After 60 minutes, aliquotes were processed for phosphorimaging. Quantitative profiles for 3 model substrates were created. The percentage of each substrate rescued by the addition of the DUBs is reported (two independent experiments). **(c)** Dose response analysis, performed as in **(b)**. The tested concentration is shown; the value in parentheses indicates the fold-increase of the tested concentration relative to the endogenous (two independent expeirments). One representative experiment is shown in Fig. S6**(c)** and **(d)** respectively. Performed as in **(b)** with the exception that ubiquitin was not added with the DUBs. Samples were processed for immunoblotting.

First, we confirmed the activity of each recombinant DUB with a UbVS-reactivity assay (Fig. S6b, c). All DUBs tested showed almost complete labelling with UbVS, with the exception of USP14 that requires activation by the proteasome(B.-H. Lee et al. 2010a; Borodovsky et al. 2001), which was absent from these in vitro assays. We then proceeded to add each recombinant DUB to extract pre-treated with UbVS/ubiquitin and monitored the stability of ^35^S-m ethionine-labeled ING2, CSNK1E, and CRIPT (Fig.3b). We chose these proteins as model substrates because they have not been reported to be regulated by DUBs and they have unrelated functions. We began by assessing all DUBs at the same concentration (800 nM), which is substantially greater than the endogenous concentration for ten of the fourteen DUBs tested (Fig. S7a). We compared the ability of each DUB to rescue degradation of the selected substrates, creating a “DUB profile” that reports the percentage of substrate rescued by each DUB (Fig. 3b). We observed that only USP7, USP15, USP25 and USP28 could rescue degradation of these substrates (Fig. 3b, Fig. S7b). Whereas USP28 and USP15 are present at low concentration in extract (4 and 20 nM respectively), USP25 was not detected in the previous study indicating that its concentration is very low, so its activity was not further evaluated. USP7 is much more abundant in extract (150 nM), suggesting that it might be able to rescue degradation of these substrates at its physiological concentration. In fact, when tested at physiological concentrations, USP7 remained capable of strongly rescuing CRIPT stability, with partial rescue of ING2 stability (Fig.3c, S7c). In contrast, USP15 and USP28 did not rescue substrate degradation when used at lower concentrations, even though these exceeded their endogenous concentrations by 5- to 15-fold, respectively (Fig. 3c, S7c). Therefore, among all the DUBs screened, USP7 seemed unique in its ability to rescue degradation of the substrates when tested at its physiological concentration. In contrast, USP4, USP9X, OTUD3, USP8, USP10, and USP21 did not rescue degradation of the three substrates, even though they were tested well above their endogenous concentrations. We were puzzled by the fact that the two known proteasome associated DUBs, UCHL5 and USP14, did not affect degradation of the substrates tested. Because they are very abundant in extract, we retested them and we tested UCHL3, another very abundant DUB in extract, at 5 times their endogenous concentration and we compared them to USP7. None of these DUBs had an effect comparable to USP7 in this assay (Fig. 3d, S7d).

Because USP7, USP15, USP25, and USP28 were each sufficient to stabilize all three substrates when used at 800 nM concentration, we wondered whether these DUBs were also capable of restoring ubiquitin availability in UbVS-treated extracts. We treated extract with 10 μM UbVS, added each of the recombinant DUBs and monitored the charging status of Ube2C-Ub thioester as a readout of ubiquitin availability. We included USP21 as a positive control, as it broadly deubiquitylates substrates *in vitro*, as well as USP5, which can regenerate free ubiquitin by acting on unanchored ubiquitin chains(Hadari et al. 1992). As expected, addition of exogenous ubiquitin fully restored charging of the Ube2C-Ub thioester (Fig.3e), as did addition of USP21, consistent with its known broad substrate specificity. USP5 did not have any effect, suggesting that free unanchored ubiquitin chains are not generated at high levels in UbVS-treated extract. Of the 4 DUBs able to rescue substrate degradation, only USP15 was able to fully restore charging of Ube2C. This finding suggests that USP7, USP25, and USP28 are capable of rescuing substrate degradation without impacting ubiquitin availability in the extract, whereas USP15 seems to have a different substrate specificity that allows it to rescue degradation and also restore ubiquitin recycling. On the other hand, despite the ability of USP21 to regenerate free ubiquitin, it is unable to rescue the degradation of any of the tested substrates.

### USP7 broadly rescues protein degradation in UbVS/ubiquitin treated extract

As USP7 seemed to be the most efficient DUB in rescuing degradation of the substrates tested, we decided to investigate how broad an effect USP7 had in rescuing proteins from degradation in extract treated with UbVS/ubiquitin. We performed a TMT-based quantitative proteomics experiment comparing protein stability in extract treated with UbVS/ubiquitin, and in extract in which USP7, USP9X, or USP4 were added at 800 nM each (Fig. 4a). We tested USP9X and USP4 as they were targets of HA-UbVS (Fig. 3a) yet were not sufficient to rescue the degradation of the panel of substrates that we tested (Fig 3b). We found that USP7 could rescue 16 of the 20 proteins degraded in UbVS/ubiquitin (Fig. 4c). However, we did not find any protein stabilized by addition of USP4 or USP9X (Fig.4b, 4c), consistent with the results of our earlier screen using the panel of ^35^S-labeled substrates. The fact that human recombinant USP7 can stabilize endogenous *Xenopus* proteins suggests again that DUB-dependent regulation of the stability of these proteins is likely evolutionarily conserved.

**Fig.4.**
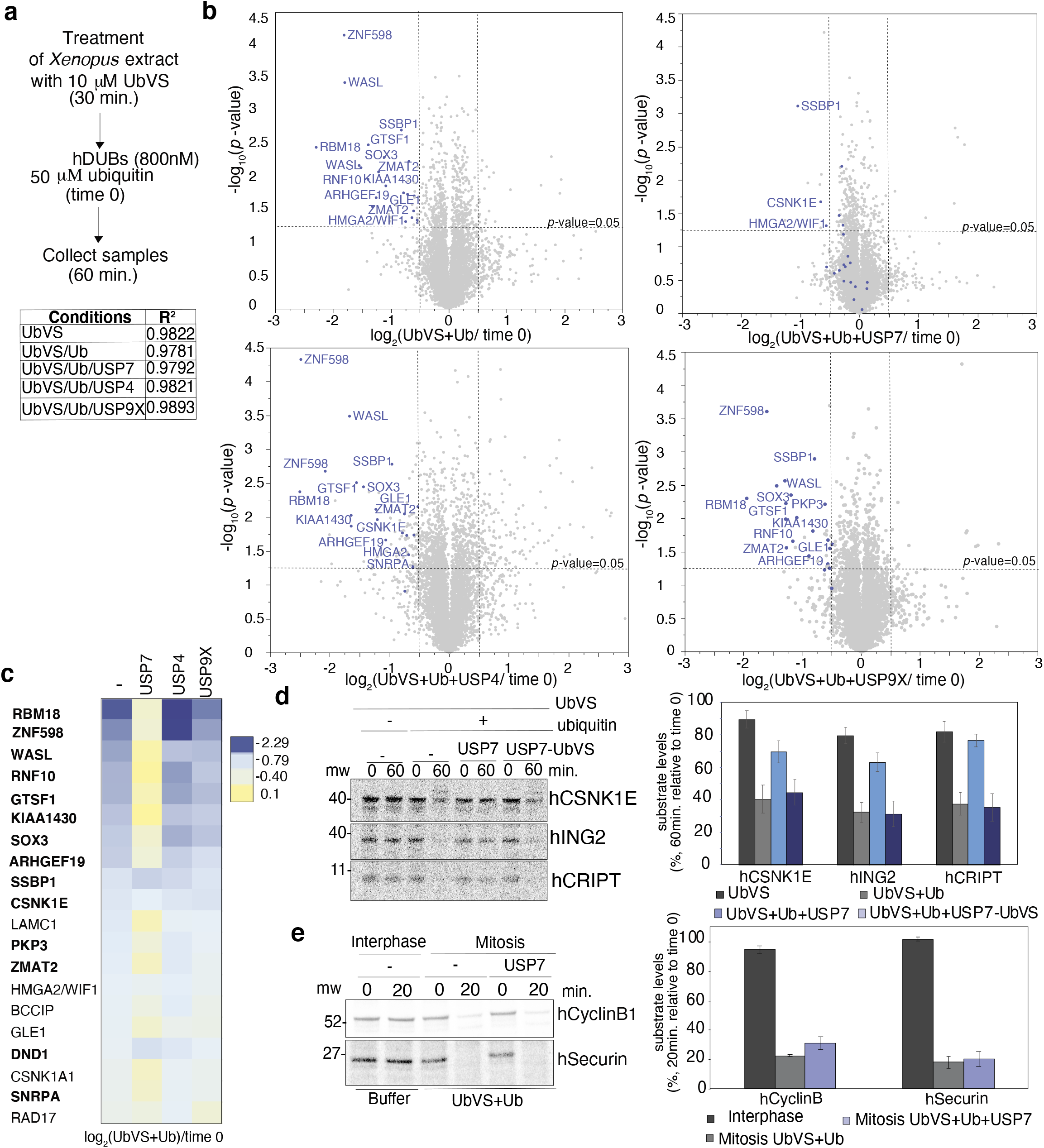
USP7 broadly rescues substrates from degradation in UbVS/ubiquitin-treated extract. **(a)** Extract was incubated with UbVS followed by addition of DUBs and ubiquitin. Samples were analyzed by SL-TMT method. Bottom: R2 of the technical replicates**(b)**Volcano plot of quantitative proteomics analysis comparing proteins detected in the different conditions. Statistical significance is plotted against the log transformed ratio of the samples collected in duplicate. Proteins significantly decreasing in UbVS/ubiquitin are in blue **(c)** Heat map of the proteins downregulated in UbVS/ubiquitin after addition of DUBs. Bold: Proteins decreasing in both experiments in Fig. 1f. **(d)** Extract was treated such as in **(a)** 35S-methionine labeled substrates were added (time 0). USP7 was pre-incubated (30 min.) with buffer or UbVS (USP7-UbVS).Right:Quantification. Error bars: SEM of three independent experiments. One representative experiment is shown. (e) 35S-methionine labeled substrates were added to interphase and mitotic extract incubated with UbVS (30 min.). USP7 and ubiquitin were added (time 0). Aliquots were collected and processed for phosphorimaging. Right: Quantification of the substrate present at 20 minutes Error bars: SEM of three independent experiments. One representative experiment is shown.

Because USP7 broadly rescued substrate degradation in UbVS/ubiquitin-treated extract (Fig. 4b, 4c) and has been reported to be associated with the proteasome(Bousquet-Dubouch et al. 2009; Besche et al. 2009), USP7 might inhibit the proteasome independent of its catalytic activity, as is known for USP14(B.-H. Lee et al. 2010a). Thus, we tested if the catalytic activity of USP7 was required for substrate rescue. Pre-incubation of USP7 with UbVS inhibited rescue of substrate degradation (Fig. 4d), confirming that deubiquitylating activity of USP7 is required for the rescue. Second, we investigated if USP7 could inhibit degradation of other known proteasome substrates. We tested if USP7 can rescue the APC substrates cyclin B1 and securin from proteasome degradation. Surprisingly, we found that USP7 was unable to rescue them from degradation (Fig. 4e), indicating that USP7 does not generally inhibit the degradation of all substrates of the UPS.

### Inhibition of Usp7 is not sufficient to induce protein instability suggesting that DUBs function redundantly to control protein stability in extract

Because USP7 could stabilize most of the proteins degraded in extracts treated with UbVS/ubiquitin, we tested if inactivation of USP7 using the recently developed specific inhibitor XL-188(Lamberto et al. 2017) was sufficient to induce degradation of the substrates that we identified. We also tested the effect of USP14 inhibition on proteome stability using the specific inhibitor IU-47(Boselli et al. 2017) alone or in combination with XL-188, since USP14 inactivation promotes degradation of some proteasome substrates in vitro and in human cells(B.-H. Lee et al. 2010b). First, we verified these compounds were able to inhibit the endogenous *Xenopus* DUBs. We pre-incubated extract with the active compounds (IU-47 and XL-188) and the respective inactive derivatives (IU-C and XL-203) for 30 minutes, and then added 10 μM HA-UbVS. As expected, only the active compounds were able to prevent HA-UbVS labelling of the specific targeted DUB (Fig. S8a). After we verified that the inhibitors were working in extract, we compared protein stability in extract pre-treated with 10 μM UbVS, 100 μM IU-47 (USP14i), 200 μM XL-188 (USP7i) or the combination of the latter inhibitors, with TMT-based quantitative proteomics (Fig. S8b). In all conditions we added ubiquitin to minimize any impact of DUB inhibition on ubiquitin recycling. In this experiment, we confirmed again that 14 of the 34 candidates (Fig. S8c) were degraded in UbVS/ubiquitin treated extract. Among these proteins, only HN1 was destabilized after specific USP7 inhibition. There were no other proteins (detected with more than one peptide) that were destabilized following specific USP7 inhibition. Inhibition of USP14 did not promote protein instability in *Xenopus* extract, as none of the proteins degraded in UbVS/ubiquitin, or any other protein, was degraded in IU-47-treated extract (Fig. S8c, d). Inhibition of both DUBs, as expected, caused the degradation of HN1 that was destabilized by the USP7 inhibitor. In addition, we observed degradation of three other proteins: DBN1, KHLC1, and Supervillin (SVIL), a known USP7 substrate(Fang and Luna 2013) (Fig.S8c, bottom). Together these findings suggest that inhibition of USP7 alone or in combination with USP14 is not sufficient to drive the broader pattern of protein instability that we observe in extracts treated with UbVS/ubiquitin. Instead, our results support a model in which multiple DUBs play redundant functions in maintaining stability of these substrates.

## Discussion

Our study offers the first broad picture of the dynamics of ubiquitylation, deubiquitylation, and protein degradation in *Xenopus* extract, a model system that has been used extensively to study the ubiquitylation and degradation of specific substrates. We provide important new insights into the relative roles of UbVS-sensitive DUBs in recycling ubiquitin and in protecting proteome stability. Identifying DUB substrates is challenging due to the fact that multiple DUBs can act redundantly on the same substrates. Here, by broadly inactivating DUBs we circumvented DUB redundancy and, using an integrated set of unbiased proteomic experiments, we identified both degradative and non-degrative targets of DUBs. Using these newly discovered set of physiological substrates whose stability is DUB-dependent, we investigated the ability of a panel specific DUBs to protect substrates from degradation. We revealed that USP7 was uniquely capable of rescuing most proteins from degradation in extract in which DUBs were broadly inhibited. However, since inhibition of USP7 alone was not sufficient to promote substrate degradation, our findings suggest that USP7 functions redundantly with other DUBs. In summary, our study provides the first comprehensive characterization of protein stability and ubiquitin dynamics in *Xenopus* extract, reveals novel DUB substrates and presents a new approach to characterize DUB specificity.

By analyzing protein abundance over time in interphase *Xenopus* extract, we observed that the flux of proteins targeted to the proteasome is low and remarkably resistant to UPS perturbation. Untreated extract showed little change in protein abundance in the absence of protein synthesis, indicating that the proteome is stable. This degree of proteome stability is consistent with the fact that the *Xenopus* egg has completed its growth and sits in a quiescent state until fertilization. Yet at the same time, ubiquitylation and deubiquitylation appear to be highly active. Our findings suggest that the robustness of protein stability to UPS perturbation arises because most ubiquitylation and deubiquitylation in unperturbed extracts occurs on non-degrative substrates. Addition of ubiquitin alone led to ubiquitin incorporation into hundreds of proteins in a manner that did not impact their stability. Furthermore, addition of UbVS alone led to alterations in protein ubiquitylation but caused no change in protein stability, instead causing a rapid depletion of free ubiquitin. This finding highlights the predominant role of UbVS-sensitive DUBs in recycling ubiquitin in extracts. Included among the non-degradative targets of UbVS-sensitive DUBs, we identified highly abundant proteins such as histones and ribosomes that may serve to sequester ubiquitin to “store” it for other purposes, such as stress resistance. Consistent with this idea, proteotoxic stresses such as heat shock or proteasome inhibition that induce a high demand for free ubiquitin, cause a redistribution of ubiquitin from histones to misfolded proteins or to proteins targeted to degradation in actively growing cells(Dantuma et al. 2006),(Rose et al. 2016)

Although most ubiquitylation in *Xenopus* extract appears to be non-degradative, our experiments also identified a set of novel degradative substrates whose stability requires ongoing deubiquitylation by UbVS-sensitive DUBs. DUB-dependent stability of these proteins was revealed only when DUBs were broadly inhibited, and available ubiquitin was restored. We confirmed DUB-dependent stability of these substrates with recombinant human proteins, demonstrating evolutionary conservation. Several substrates are known to be regulated by specific DUBs, including MARCH7, BIRC3, STAM and NFX1 (Nathan et al. 2008),(Mei et al. 2011),(Chen et al. 2019),(Cai et al. 2015). Furthermore, PIM3 and WASL have not been connected previously to DUBs, but are similar in sequence to well-known DUB substrates (PIM2(Kategaya et al. 2017) and WASH(Hao et al. 2015) respectively). Together, these findings validate that our approach can identify established DUB substrates. Still, the vast majority of the proteins we identified are not known DUB substrates, demonstrating the novelty of our findings. Our candidate substrates were enriched for ubiquitin ligases (n=12), extending our understanding of the importance of DUBs in counteracting their auto-ubiquitylation and consequent degradation(W. Kim et al. 2011),(Ventii and Wilkinson 2008). Makorin Ring Finger protein 1 (MKRN1) ubiquitylates p53 and p21 targeting them to proteasomal degradation(E.-W. Lee et al. 2009). Yet despite intensive study, a role for DUBs in controlling MKRN1 stability has not been described. Our data suggest that DUB-dependent stability of MKRN1 could be an important control mechanism, as it is for MDM2, another p53 ubiquitin ligase(Ranaweera and Yang 2013).

Beyond ubiquitin ligases, we identified substrates with a broad range of functions. We found multiple transcription factors (n=7) and proteins involved in RNA metabolism (n=12). A number of substrates, such as SPRTN, ETAA1, and the Casein Kinases ε and δ, have been extensively studied, so it is surprising that DUB-dependent control of their stability has not yet been reported. SPRTN and ETAA1 are both involved in DNA replication/damage and are essential for maintaining genome stability(Vaz et al. 2016),(Haahr et al. 2016). Casein Kinases ε and δ are versatile proteins that participate in multiple processes such as cell cycle control, spindle organization, and circadian rhythm(Schittek and Sinnberg 2014). Our study suggests that DUBs could modulate their degradation by the UPS as mechanism to regulate their activity.

This study also describes a new approach for studying the specificity and activity of DUBs in a system in which physiological rates of substrate ubiquitylation are maintained. Our approach measures effects on degradation rather than deubiquitylation, providing a new way of assessing the ability of DUBs to counter proteasome-mediated degradation. The most unexpected finding to emerge from this analysis was the ability of USP7 to rescue a large number of substrates from degradation. Despite this broad activity, USP7 was unable to rescue APC/C substrates from degradation, suggesting that USP7 is not a general inhibitor of proteasome activity. This result was not due to a lack of activity of USP7 in mitosis because we verified that USP7 was still fully active in mitosis (data not shown). Furthermore, even though USP7 was able to broadly rescue substrates from degradation, USP7 was unable to rescue ubiquitin depletion in UbVS-treated extract. Together these findings indicate that USP7 has a defined substrate specificity that seems targeted towards degradative versus non-degradative substrates. Interestingly, we also observed exactly the opposite pattern of specificity with USP21(Yu Ye et al. 2011), which was able to rescue ubiquitin availability but was unable to rescue degradation of our panel of substrates.

What explains the broad ability of USP7 to antagonize proteasomal degradation? USP7 has been highly studied and has attracted attention as a pharmacological target, as it regulates the stability of the tumor suppressor p53 and its regulator MDM2(Sheng et al. 2006). Among the DUBs we tested, USP7 has the greatest number of published substrates, most of which are targeted to the proteasome(R. Q. Kim and Sixma 2017). Among the substrates that we identified, a few such as MARCH7, SVIL, NEK2A, and TRIP12 are known targets of USP7(Nathan et al. 2008),(Fang and Luna 2013),(Franqui-Machin et al. 2018),(Cai et al. 2015) whereas two others, PIM3 and WASL, are likely USP7 targets since it deubiquitylates related proteins (PIM2(Kategaya et al. 2017) and WASH(Hao et al. 2015) respectively) in human cells. USP7 has been reported to associate with the proteasome(Bousquet-Dubouch et al. 2009),(Besche et al. 2009), where it could intercept proteins that are targeted for degradation. However, the physiological significance and mechanism of this association has not been carefully studied. USP7 may have a broad ability to rescue substrates from degradation because it directly binds a wide range of proteins through recognition of common degenerate motifs (P/AxxS and Kxxx/KxK) that are found in many proteins(R. Q. Kim and Sixma 2017). Another plausible explanation is that USP7 is kinetically sufficient to counteract the rates of ubiquitylation of ubiquitin ligases that target substrates for degradation. This idea is supported by USP7 being the most active DUB in Ub-AMC assays among a panel of 12 cysteine DUBs tested(Faesen et al. 2011).

Our study also highlights the important impact and challenge of functional redundancy when studying DUB activity and specificity. In the *Xenopus* system, we observed that inhibition of USP7 alone was not sufficient to induce degradation of substrates, despite USP7’s ability to broadly rescue substrates from degradation. This finding supports the notion that DUBs function redundantly to maintain the stability of the proteins in this system. Because degradation is an irreversible step in the protein lifecycle, DUB redundancy may help set a higher threshold for ubiquitylation required for degradation of a protein, beyond what is required for direct recognition by the proteasome. On the other hand, DUB redundancy poses a serious technical challenge to fully understand the role of DUBs and identify their substrates, especially using genetic approaches. Our work highlights how unmasking DUB redundancy it is key not only for discovery of novel DUB substrates but also to fully understand functional specificity of these important enzymes.

## Materials and Methods

### Gene nomenclature

The human gene symbols have been used for consistency and simplicity in the manuscript.

### Reagents

Commercial antibodies used for Western blotting analysis were as follow: anti-Ube2C (A-650, Boston Biochem), anti-ubiquitin (P4D1; sc-8017; Santa Cruz Biotechnology), anti-HA-peroxidase (3F10; Roche), anti-USP7 (A300-033A; Bethyl). Secondary antibodies used included anti-rabbit IgG-HRP (NA934V) and anti-mouse IgG-HRP (NA931V) from GE Healthcare. Chemicals used in this study included Ubiquitin vinyl sulfone (U-202), MG262 (I-120), Ubiquitin vinyl sulfone tagged with HA (U-212) and HA-ubiquitin (U-110) purchased from Boston Biochem. For immunoprecipitation anti-HA-Agarose beads (A2095) from Sigma were used. For the diGly experiment immunoaffinity beads from Cell Signaling were used (5562). Ubiquitin (U-100H) and recombinant DUBs were provided by Boston Biochem (E-519, USP7; E-520, USP8; E-320, USP5; E-325, UCHL3; E-325, UCHL5; E-546, USP25; E-552, USP9x; E-570, USP28; E-574, OTUD3; E-592, USP10; E-594, USP15; E-596, USP4).

### Preparation of *Xenopus laevis* egg extract

Interphase extract was prepared as previously described^(Murray AW 1991)^ but using 2 μg/ml calcium ionophore (A23187, Calbiochem) for egg activation. Entry into mitosis was induced by addition of 1 μM non-degradable cyclin B (MBP-Δ90) as previously described^(Zeng et al. 2010)^.

### HA-UbVS immunoprecipitation

Following treatment with 10 μM HA-UbVS (10 minutes at 24°C), *Xenopus* extract was diluted three times with XB buffer (10 mM potassium HEPES pH 7.7, 500 mM KCl, 0.1 mM CaCl2, 1 mM MgCl2, 0.5% NP40) and incubated with anti-HA beads or empty beads at 4°C (1h and 30 minutes). At the end of the incubation, beads were washed three times with XB buffer, SDS sample buffer was added and samples were subjected to SDS gel electrophoresis. Samples were processed according to the GeLC-MS/MS strategy(Paulo 2016).

### HA-UbVS immunodepletion

*Xenopus* extract was pre-treated with 10 μM HA-UbVS (10 minutes at 24°C) diluted three times with XB buffer and incubated with anti-HA beads or empty beads overnight at 4°C. Supernatants from the beads were collected and treated for Mass Spectrometry analysis by SL-TMT method (described below).

### HA-ubiquitin immunoprecipitation

Following treatment with 50 μM HA-ubiquitin (30 minutes at 24°C), *Xenopus* extract was diluted three times with XB buffer (10 mM potassium HEPES pH 7.7, 500 mM KCl, 0.1 mM CaCl2, 1 mM MgCl2, 0.5% NP40, Pierce proteases inhibitor tablet and NEM 5mM) and incubated with anti-HA beads or empty beads at 4°C (2h). At the end of the incubation, beads were washed four times with XB buffer and elute with the HA peptide (two times). Samples were processed for Mass Spectrometry analysis by SL-TMT method (described below).

### Immunoprecipitation of diGly-containing peptides

Dried peptides (2 mg of proteins) were resuspended in IAP buffer [50 mM MOPS (pH 7.2), 10 mM sodium phosphate and 50 mM NaCl] and centrifuged at top speed (5 min). After that the supernatants were added to the diGly resin (Cell Signaling Technology) and incubated for 2 hr at 4°C. After that, beads were washed three times with ice-cold IAP buffer and twice with PBS. The diGly peptides were eluted twice with 0.15% TFA, desalted using homemade StageTips and dried via vacuum centrifugation. Peptides were immunoprecipitated twice. Samples were processed for Mass Spectrometry analysis by SL-TMT method (described below).

### Degradation of ^35^S-methionine labeled substrates

Extract was pre-treated with UbVS (10 μM) for 10 min at 24°C before addition of ubiquitin (50 μM) and substrates. The preincubation time with UbVS was extended to 30 minutes when recombinant DUBs were added to the extract. Pre-treatment of extract with MG262 (200 μM) or specific DUB inhibitors was done at 24°C for 30 minutes. Substrates were expressed and labeled using ^35^S-methionine (Perkin Elmer, NEG709A500UC) with the TNT kit (Promega: L1770). Each substrate was amplified with primers by PCR to allow T7-dependent transcription of the PCR product or transcribed directly if plasmids contained a T7 promoter. The translation reaction mix was added to the *Xenopus* extract at 8% final volume. Samples of the reactions were collected at the indicated time, quenched with sodium dodecyl sulfate (SDS) sample buffer, and processed for SDS gel electrophoresis and phosphor imaging (Bio-Rad PMI). Quantification was performed using Quantity One software (Bio-Rad).

### UbVS labeling of recombinant DUBs to verify their activity

Each DUB (1μM) was incubated with saturating amount of UbVS or HA-UbVS as indicated (1 hour at 30°C). Reactions were stopped with addition of sodium dodecyl sulfate (SDS) sample buffer and run on SDS-PAGE. After SDS-PAGE, the gel was stained with Coomassie Brilliant Blue.

### Streamlined Tandem Mass Tag Protocol

#### Peptide extraction and digestion

The TMT labeling protocol and mass spectrometry analysis were based on the SL-TMT sample preparation strategy(Navarrete-Perea et al. 2018). *Xenopus* extract was collected, resuspended in the appropriate buffer (1% SDS, 5 mM DTT, 50 mM Tris pH 8.8 and Pierce protease inhibitor tablet) and flash frozen in liquid nitrogen. Methanol–chloroform precipitation was performed. Four volumes of methanol were added to each sample and briefly vortexed. One volume chloroform was added to the samples and vortexed again. Lastly, three volumes water was added and vortexed. The samples were centrifuged (5 minutes, 14000 RPM) and subsequently washed twice with cold methanol. Samples were resuspended in 200 mM EPPS, pH 8.5 and digested overnight at 24°C with Lys-C protease (Wako Chemicals). Later samples were incubated for 6 hours at 37°C for digestion by trypsin protease (Pierce Biotechnology).

#### Isobaric labeling and fractionation

Tandem mass tag (TMT) isobaric reagents (Thermo Fisher Scientific) were resuspended in anhydrous acetonitrile (final concentration of 20 μg/μL). 10 μL of the labeling reagents plus 30 μL of acetonitrile was added to the peptides obtained by the previous digestions (~100 μg). After incubation at room temperature (90 minutes), the reaction was quenched using hydroxylamine to a final concentration of 0.3% (v/v). The TMT-labeled samples were pooled at a 1:1 ratio across all the samples. Fractions were fractionated off-line using basic pH reversed phase chromatography (BPRP). Fractions were pooled into 6 or 12 super-fractions which were acidified with formic acid to a final concentration of 1%. The pooled peptides were desalted using homemade StageTip, dried with vacuum centrifugation, and reconstituted in 5% acetonitrile and 5% formic acid for LC-MS/MS processing.

#### Peptide detection, identification and quantification

All samples were analyzed on an Orbitrap Fusion mass spectrometer (Thermo Fisher Scientific) coupled to a Proxeon EASY-nLC 1000 liquid chromatography (LC) pump (Thermo Fisher Scientific). Peptides were separated on a column packed with 35 cm of Accucore C18 resin (2.6 μm, 150 Å, Thermo Fisher Scientific). The column had a 100 μm inner diameter microcapillary. For each experiment, 2 μg of peptides were loaded onto this column. Peptides were separated, using a flow rate of 450 nL/min., with a 150-minute gradient of 3 to 25% acetonitrile in 0.125% formic acid. Each analysis used an MS3-based TMT method, which it is known to reduce ion interference if compared to MS2 quantification. The scan sequence starts with an MS1 spectrum (Orbitrap analysis, resolution 120,000, 400-1400 Th, automatic gain control (AGC) target 5E5, maximum injection time 100 ms). For subsequent MS2/MS3 analysis, only the top 10 precursors were selected. MS2 analysis included: collision-induced dissociation (CID), quadrupole ion trap analysis, automatic gain control (AGC) 2E4, NCE (normalized collision energy) 35, q-value 0.25, maximum injection time 120 ms), and isolation window at 0.7. After we acquire each MS2 spectrum, we collected an MS3 spectrum in which multiple MS2 fragment ions were captured in the MS3 precursor population with isolation waveforms using multiple frequency notches. MS3 precursors were fragmented by HCD and analyzed using the Orbitrap (NCE 65, AGC 1.5E5, maximum injection time 150 ms, resolution was 50,000 at 400 Th). For MS3 analysis, we used charge state-dependent isolation windows: For charge state z=2, the isolation window was set at 1.3 Th, for z=3 at 1 Th, for z=4 at 0.8 Th, and for z=5 at 0.7 Th. Collected Spectra were processed using a Sequest-based software pipeline. Spectra were converted to mzXML using MS Convert (Adusumilli and Mallick 2017). Database searching included all the entries from the PHROG databas. This database includes many lower abundant proteins and multiple splice isoforms (not present in other databases). In the original study, around 11,000 proteins were identified(Wühr et al. 2014). Thus, our study (with ~8000 proteins) represents an acceptable coverage of the *Xenopus* proteome. This database was concatenated with one composed of the sequences in the reversed order. Searches were performed using a 50 Th precursor ion tolerance and the product ion tolerance was set to 0.9 Da. Oxidation of methionine residues (+15.995 Da) and, where indicated. Peptide-spectrum matches (PSMs) were adjusted to a 1% false discovery rate (FDR). PSM filtering was performed using a linear discriminant analysis, as described previously and assembled to a final protein-level FDR of 1%. Proteins were quantified by summing reporter ion counts across all matching PSMs(McAlister et al. 2012). Reporter ion intensities were adjusted to correct for the isotopic impurities of the different TMT reagents according to manufacturer specifications. The peptides signal-to-noise (S/N) measurements assigned to each protein were summed and normalized so that the sum of the signal for all proteins in each channel was equivalent, thereby accounting for equal protein loading. Lastly, each protein was scaled such that the summed signal-to-noise for that protein across all channels was 100, thereby generating a relative abundance (RA) measurement.

### TMT Mass-spectrometry analysis

A two-sided Student’s t-test was used as a measure of statistical confidence of the observed log_2_ fold change. Selected candidates met both thresholds (Fold Change ≤ −1.5 and *p*-value<0.05) in the experiments. For Figure 1g, the candidates were included if at least one peptide was identified in both of the experiments. For subsequent figures, where a single experiment was performed, candidates were included only if they were detected with at least 2 different peptides. When multiple isoforms of the same protein decreased in UbVS/ubiquitin, only the isoform with more peptides was selected (for simplicity isoforms are not indicated in the figures).

## Acknowledgments

We gratefully acknowledge W. Harper for the gift of plasmids from the ORFeome collection, S. Buhrlage for the gift of XL-188 and XL-203 and D. Finley for the gift of IU-C, IU-47 and USP21. We thank Sandhya Manohar for the help in preparing the *Xenopus* egg extract. We thank W. Harper for helpful discussions. This work was supported by NIH grants R01 GM132129 (J.A.P.), R01 GM67945 (S.P.G) and R35 GM127032 (R.W.K.).

## Author contributions

V.R. designed, performed, and analyzed all the experiments

V.R. prepared all samples for mass spectrometry with assistance from J.P.

J.P., J.C. and S.P.G. performed mass spectrometric analysis and provide reagents

B.B. provided advice on DUB selection and DUB assays

R.W.K. assisted with experimental design

V.R. and R.W.K. conceived of the project and wrote the manuscript with input from the other authors.

## Competing interests

The authors declare no competing interests

## Data availability

The mass spectrometry proteomics data have been deposited to the ProteomeXchange Consortium via the PRIDE partner repository with the dataset identifier ….

---

**Fig. S1.**
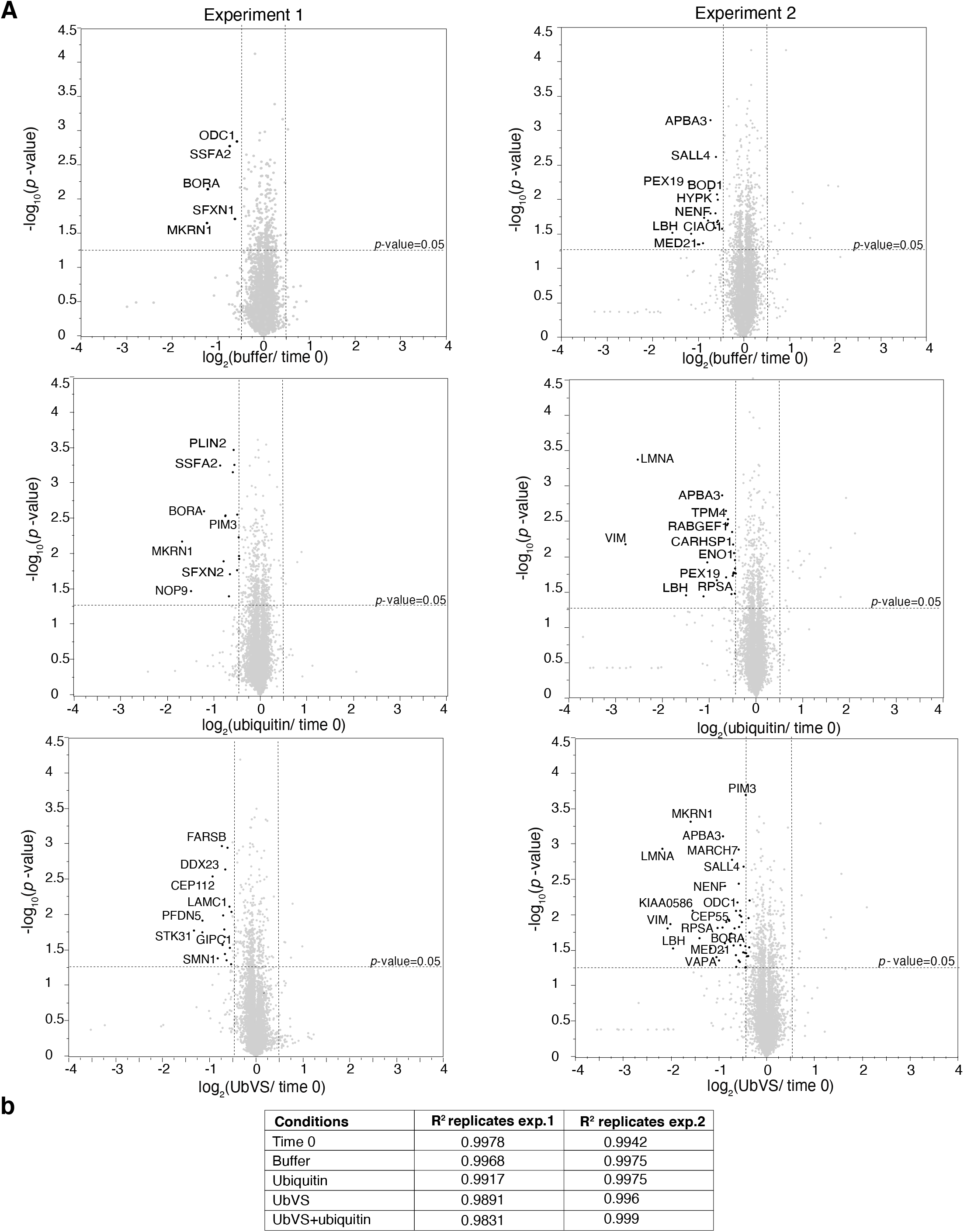
Analysis of the Xenopus proteome over time, after ubiquitin and UbVS addition. **(a)** Volcano plot of the 2 proteomic experiments comparing protein abundance between the indicated conditions (60 minutes) and time 0 as in Fig.1f. Statistical significance (−log_10_ *p-value*) is plotted against fold change (average log2 ratio). Proteins significantly decreasing in the indicated conditions are in black. (**b**) R2 of the technical replicates are reported.

**Fig. S2.**
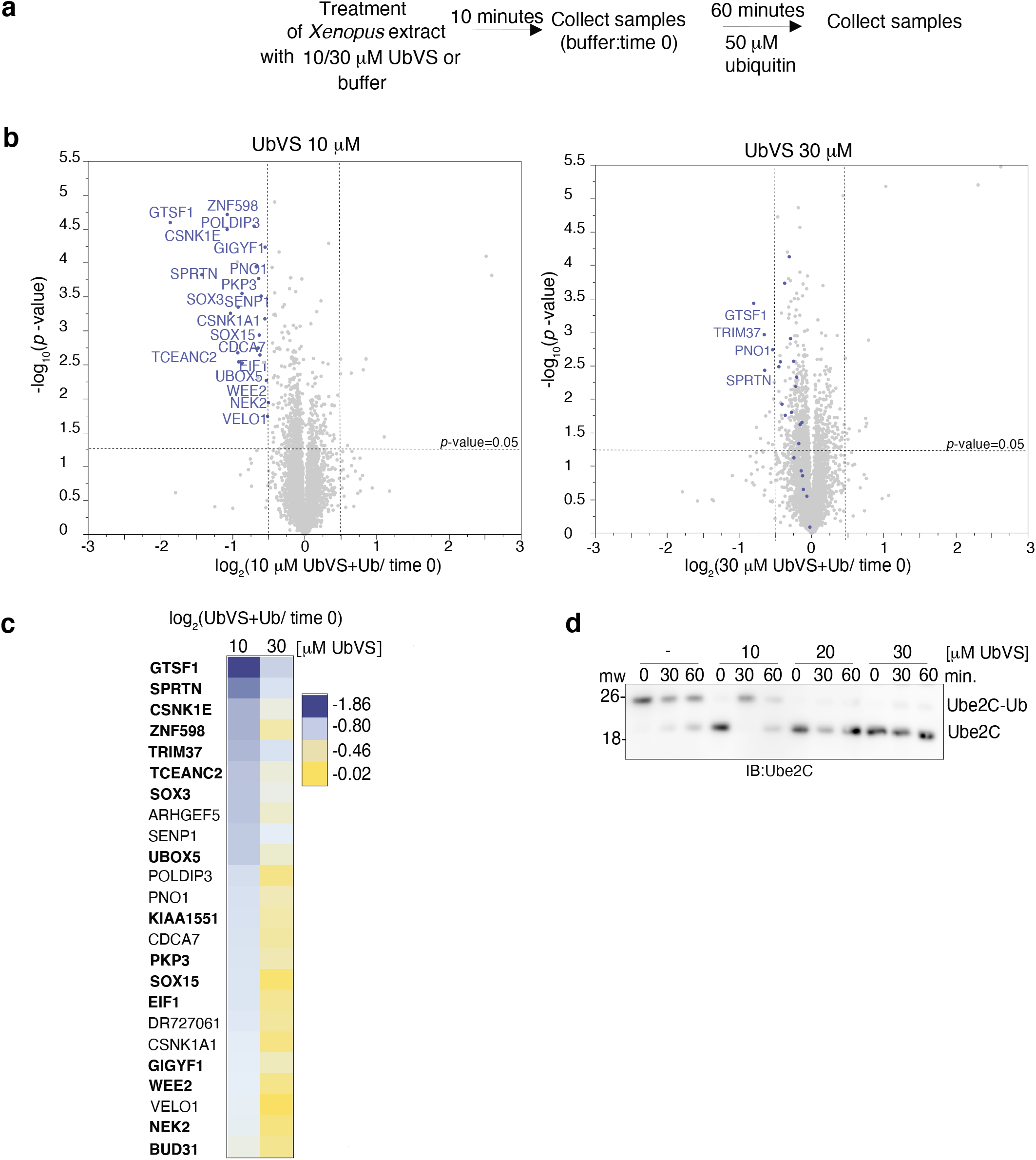
Increasing UbVS concentration does not enhance protein degradation in extract. **(a)** Extract was incubated with buffer, 10 or 30 μM UbVS (10 minutes) and 50 μM of ubiquitin was added (time 0). Samples were collected in triplicate and processed for SL-TMT method analysis **(b)** Volcano plot of quantitative proteomics analysis. Statistical significance (−log_10_ *p-value*) is plotted against ratio (average log_2_ ratio). Blue dots: proteins significantly decreasing in 10 μM UbVS/ubiquitin (Fig.1g) **(c)** Log2 ratio heat map compares the proteins decreasing in **(b)**. Proteins decreasing in the experiments in Fig.1f are in bold. **(d)** Extract was treated with UbVS (10 minutes), after which 50 μM of ubiquitin was added (time 0). Aliquots were subjected to immunoblotting.

**Fig.S3.**
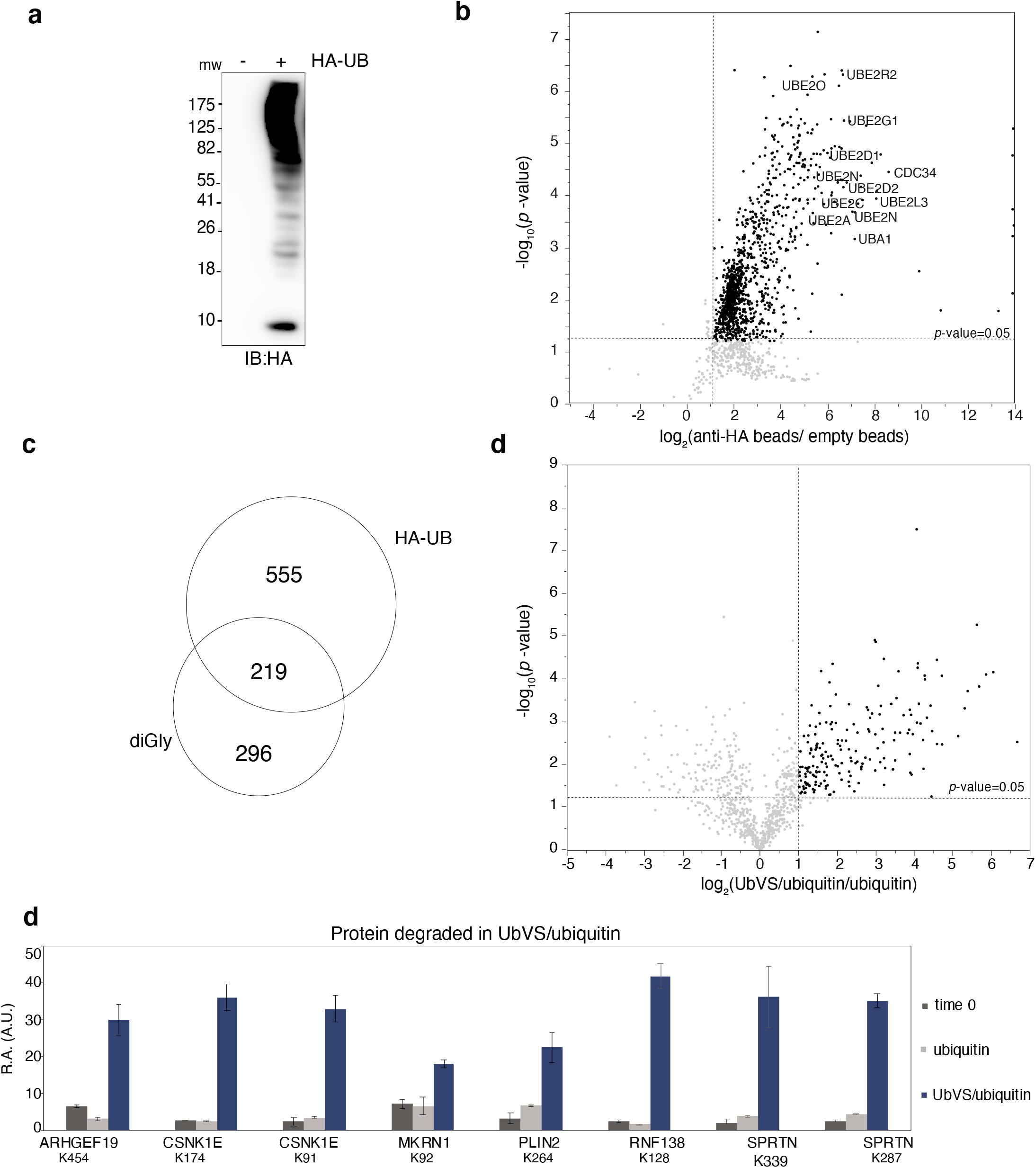
Identification of non-degradative substrate-ubiquitin conjugates. **(a)** Extract was treated with 50 μM HA-ubiquitin (HA-UB) for 30 minutes and subjected to immunoblotting (IB) **(b)** After incubation with HA-ubiquitin, extract was subjected to anti-HA pull-down and analyzed by TMT-mass spectrometry. Volcano plot comparing abundance of protein bound to anti-HA beads relative to empty beads. Statistical significance (−log_10_ *p-value*) is plotted against ratio (average log2). Samples were collected in technical triplicates. Proteins significantly increasing are in black. Abundant UPS components are labeled **(c)** Overlap between the HA-ubiquitin pull down and the diGly enrichment **(d)**Volcano plot of the TMT diGly remnant analysis comparing changes in ubiquitination sites detected after addition UbVS/ubiquitin to the changes after addition of only ubiquitin (30 min.).Samples were collected in duplicate.Statistical significance is plotted against ratio (average log2). Proteins significantly increasing are in black. **(e)** Ubiquitination sites of the proteins destabilized in UbVS/ubiquitin (Fig.1f) are shown. The graph shows the relative amount (R.A.) of ubiquitination in the conditions tested (average between duplicates). A.U. Arbitrary unit. Error bars: standard deviation (N=2)

**Fig. S4.**
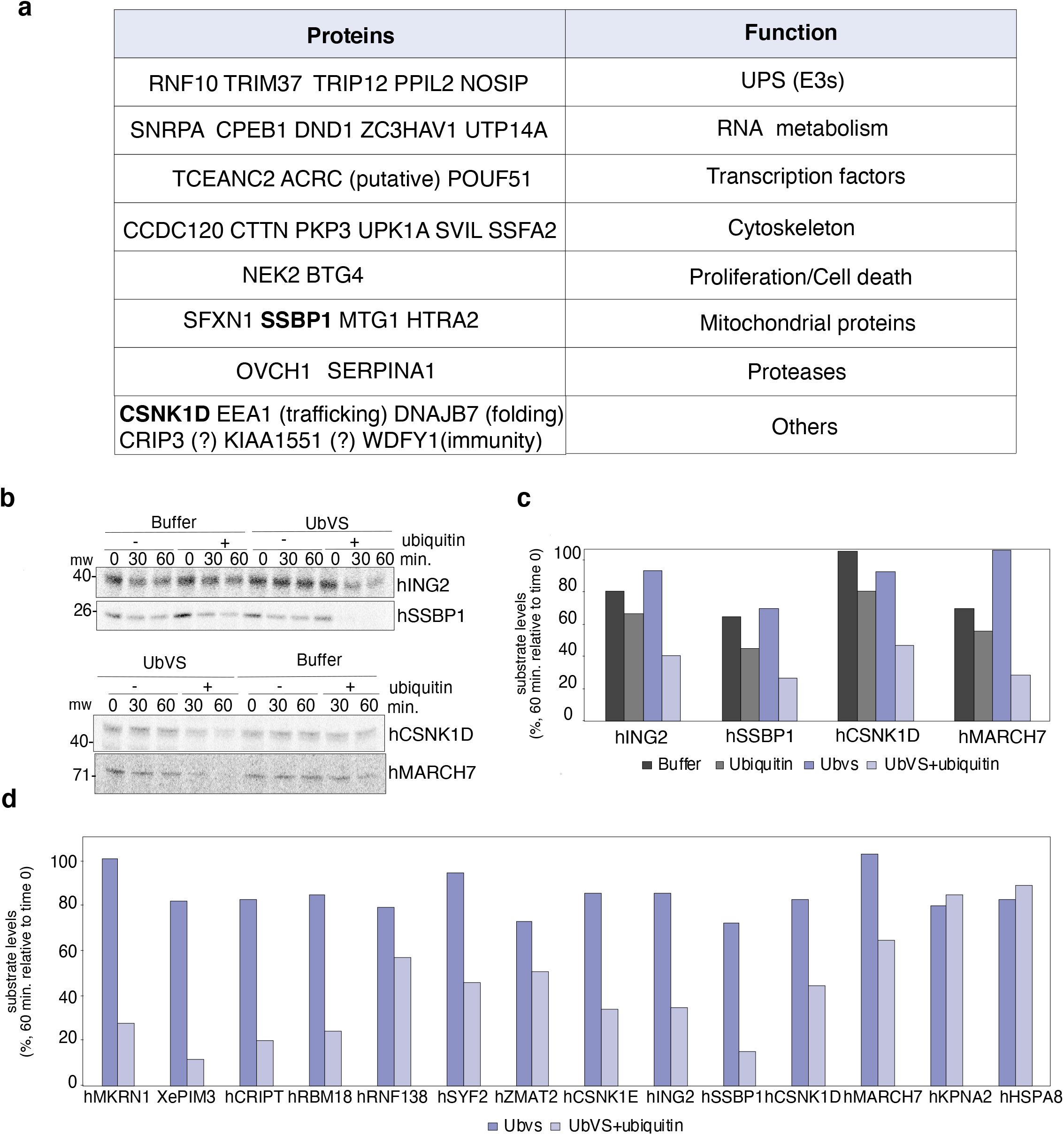
Validation of the substrates protected from degradation by the UbVS-sensitive DUBs. **(a)** Functions of the 5% of proteins whose abundance decreased most in UbVS/ubiquitin (FDR 1%) in both the experiments (Fig.1f). The 34 proteins included in the more selective threshold are not shown. In bold, proteins validated with independent experiments. **(b)** The indicated proteins were expressed and labeled as described previously and added to extract pre-treated as indicated. Levels of the proteins were assessed by SDS-PAGE and phosphorimaging **(c)** Quantification of the substrate levels (60 minutes) of the experiment in **(b)**. Substrates have been validated with 2 independent experiments. One representative experiment is shown. **(d)** Quantification of the second set of independent experiments in **(c)** and in Fig.2.

**Fig. S5.**
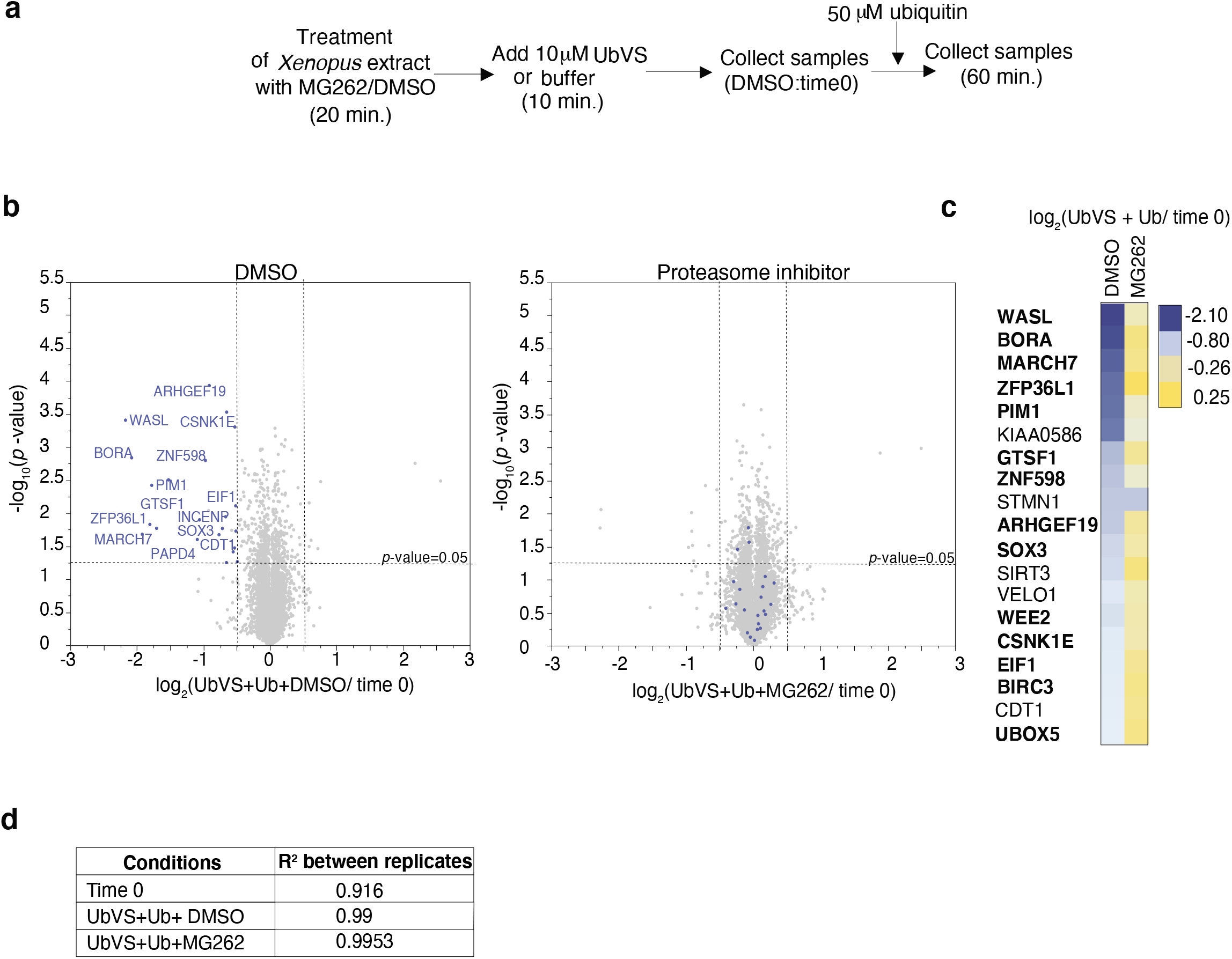
Degradation of proteins in UbVS/ubiquitin-treated extract is proteasome-dependent. **(a)** Extract was incubated with 200 μM MG262 or DMSO (20 minutes), after which 10 μM UbVS was added (10 minutes), followed by addition of 50 μM of ubiquitin (time 0). Samples were collected in duplicate (time 0 and 60 minutes) and processed for mass spectrometry analysis by the SL-TMT method **(b)** Volcano plot of quantitative proteomics analysis comparing the proteins detected in UbVS/ubiquitin in presence of DMSO or MG262 at 60 minutes with the proteins detected at time 0. Statistical significance (−log_10_ *p-value*) is plotted against ratio (average log2 ratio). Blue dots represent the proteins significantly decreasing in UbVS/ubiquitin and ubiquitin are labeled **(c)** Log2 ratio heat map of the proteins significantly downregulated in UbVS/ubiquitin. Proteins decreasing in the experiments in Fig.1G are in bold. **(d)** R2 of the technical replicates of each condition.

**Fig S6.**
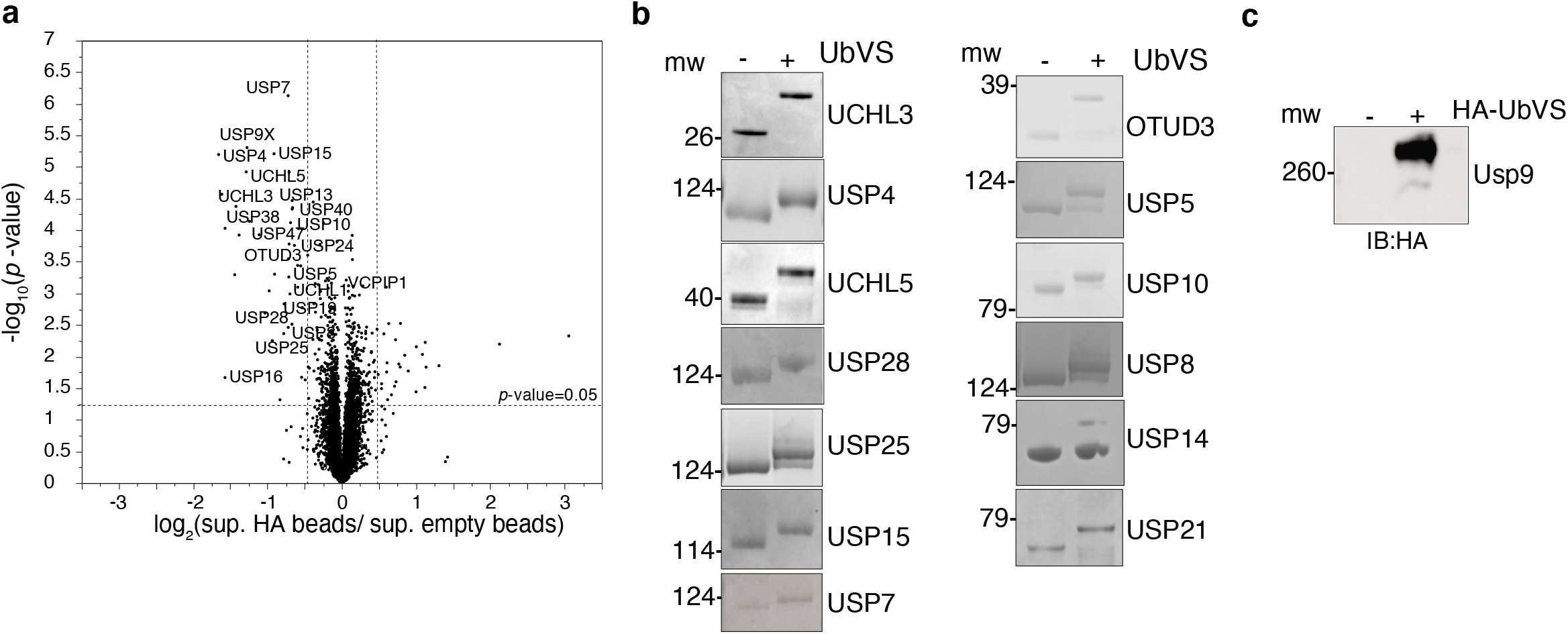
Identification of the fraction of each DUB depleted by HA-UbVS and validation of the catalytic activity of recombinant DUBs. **(a)** Volcano plot of quantitative proteomics analysis comparing the proteins detected in the supernatant of the empty beads with the proteins detected in the supernatant of the anti-HA beads. Statistical significance (−log_10_ *p-value*) is plotted against ratio (average log_2_ ratio). DUBs are labeled **(b)** Indicated DUBs were incubated with saturating amount of UbVS or buffer (30 minutes). Reactions were stopped with SDS sample buffer, run on SDS-PAGE and the gel was stained with Comassie Blue. **(c)** USP9X was incubated with HA-UbVS and its activity was examined by Immunoblotting.

**Fig.S7.**
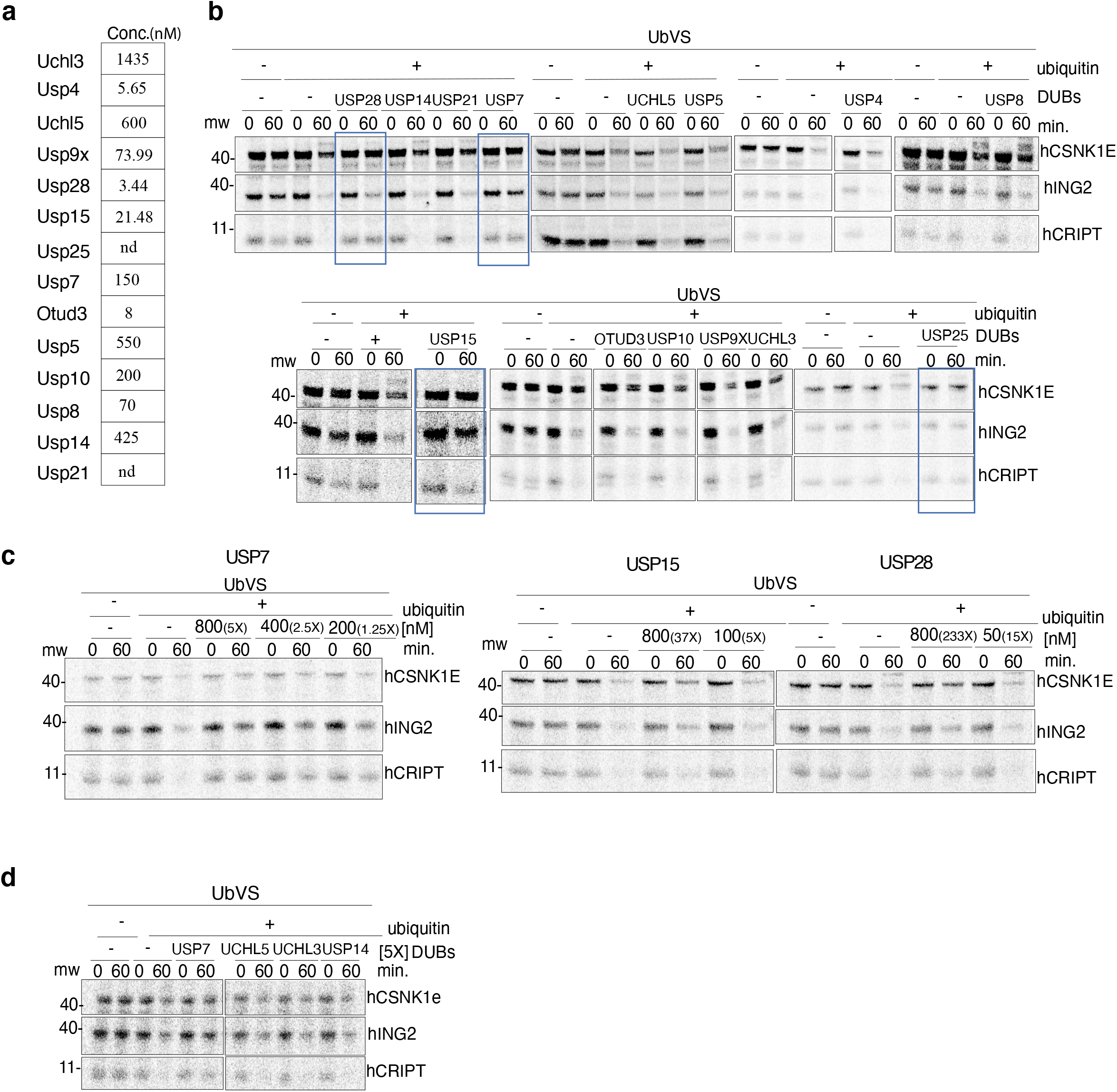
Experiments used to create the quantitative DUB activity profile (Fig.3). **(a)**Estimated endogenous concentration of the DUBs tested (Fig.3a). One of the representative experiments reported in the quantitative profile in Fig.3b is shown in **(b)**, in Fig.3c is shown in **(c)** and in Fig.3d is shown in **(d)**.The blue rectangles in **(a)** indicate the four DUBs rescuing all the three substrates tested.

**Fig.S8.**
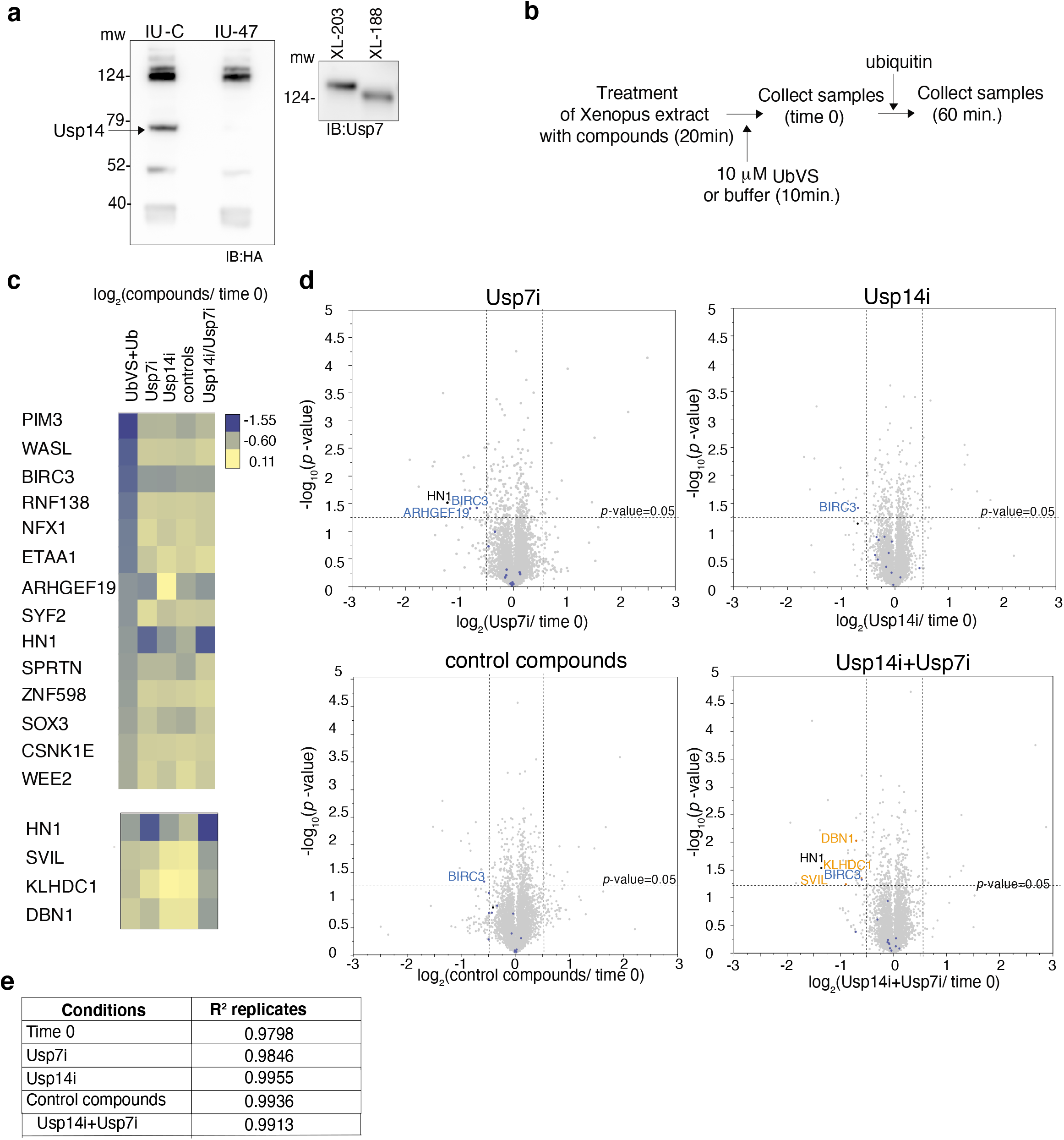
Usp7 or Usp14 inhibition does not promote broad protein instability. **(a)** Extract was incubated with the indicated compounds (20 minutes), followed by HA-UbVS addition (20 minutes). Aliquots were analyzed by immunoblotting. IU-47 and IU-C: Usp14 inhibitor (Usp14i) and control compound. XL-188 and XL-203: Usp7 inhibitor (Usp7i) and control compound. **(b)** Extract was incubated with DMSO or compounds. UbVS was added for 10 minutes(time 0) to the samples with DMSO. Ubiquitin was added to all samples, samples were collected in duplicate and analyzed by the SL-TMT method. **(c)** Top: heat map compares the effect of the specific DUB inhibitors on the proteins significantly decreasing in UbVS/ubiquitin in this experiment and in the experiments in Fig. 1F. Bottom: heat map of the proteins affected by the specific DUB inhibitors. **(d)** Volcano plot of quantitative proteomicsanalysis. Statistical significance (−log_10_ *p-value*) is plotted against ratio (average log_2_ ratio).Blue dots: proteins decreasing in UbVS/ubiquitin; black dots:proteins decreasing in UbVS/ubiquitin and in presence of Usp7i; orange dots: proteins decreasing in presence of both the inhibitors. **(e)** R2 of the technical replicates

**Table S1:**
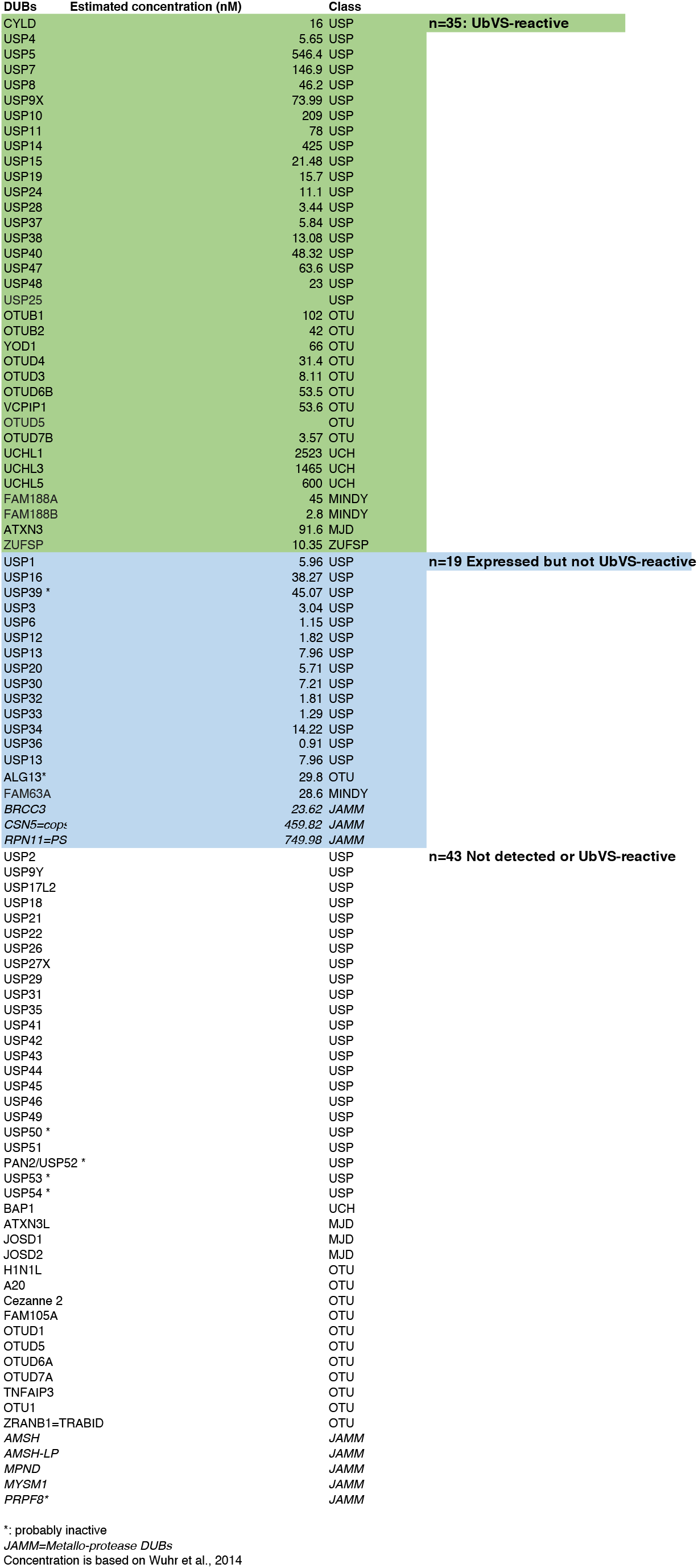
List of the Xenopus DUBs.

**Table S2:**
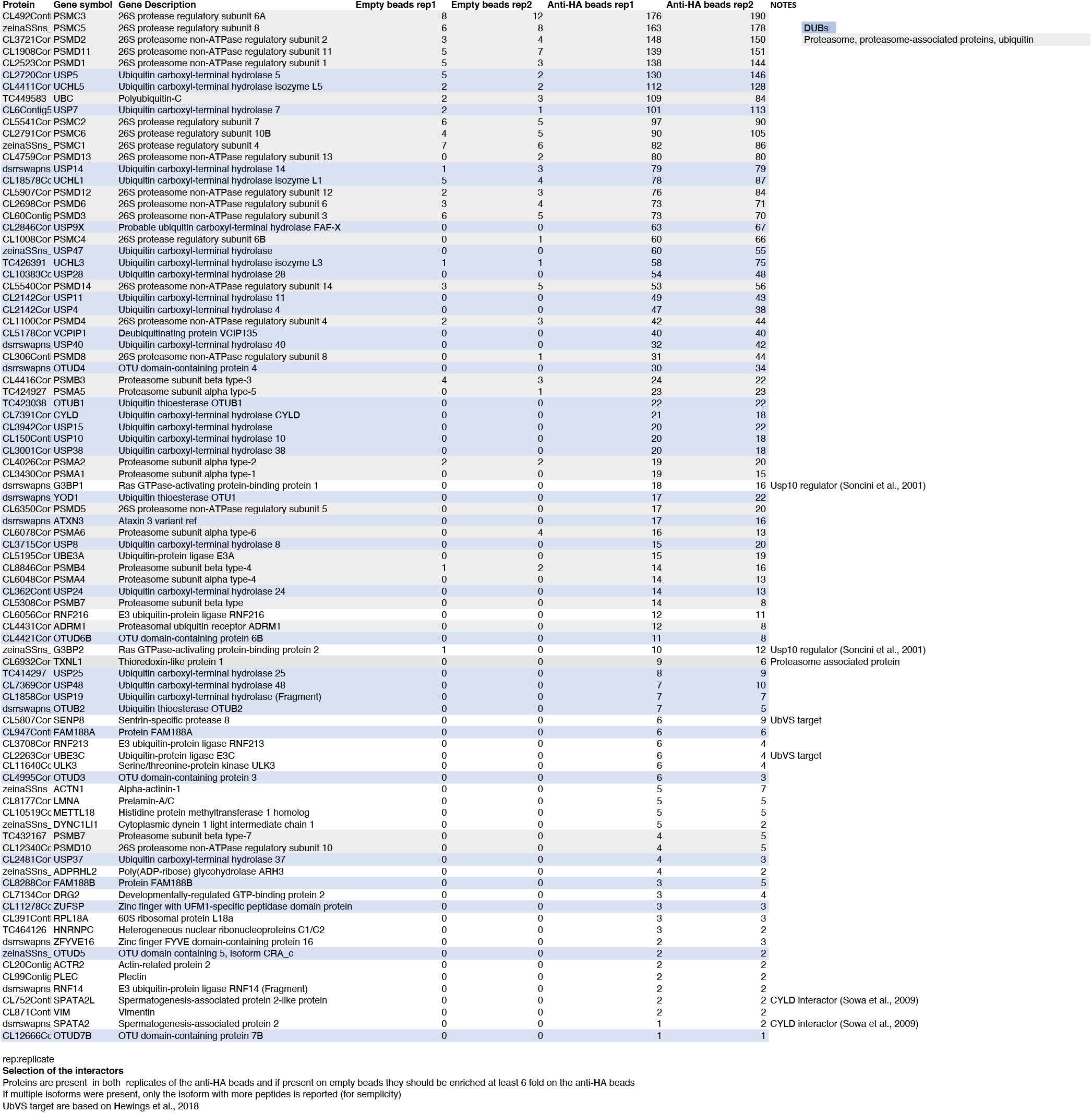
HA-UbVS interactors.

**Table S3:**
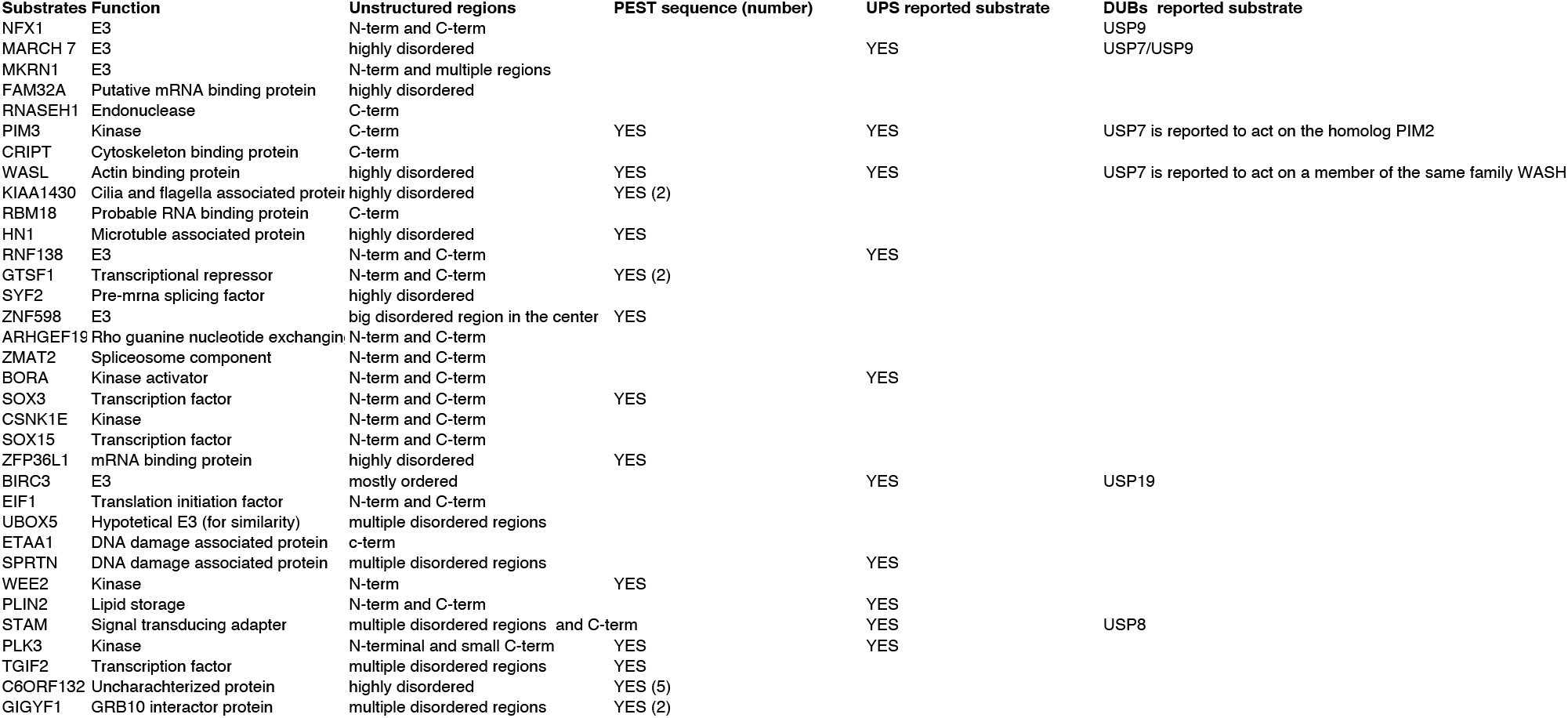
UbVS-sensitive proteolitic DUB substrates.

## Notes

### Competing Interest Statement

The authors have declared no competing interest.

